# Molecular Specification of Claustro-Amygdalar and Paleocortical Neurons and Connectivity

**DOI:** 10.1101/2024.10.20.619229

**Authors:** Navjot Kaur, Rothem Kovner, Forrest O. Gulden, Mihovil Pletikos, David Andrijevic, Tianjia Zhu, John Silbereis, Shaojie Ma, Nikkita Salla, Xabier de Martin, Thomas S. Klarić, Yuting Liu, Akemi Shibata, Daniel Franjic, Hyesun Cho, Matthew Yuen, Ipsita Chatterjee, Megan Burke, Devippriya Esakkimuthu, Markus Moser, Gabriel Santpere, Yann S. Mineur, Kartik Pattabiraman, Marina R. Picciotto, Hao Huang, Nenad Sestan

## Abstract

The ventropallial excitatory neurons (ExNs) in the claustro-amygdalar complex and piriform cortex (PIR; part of paleocortex) form crucial reciprocal connections with the prefrontal cortex (PFC), integrating cognitive and sensory information that result in adaptive behaviors. Early-life disruptions in these circuits are linked to neuropsychiatric disorders, highlighting the importance of understanding their development. Our study uncovers that transcription factors SOX4, SOX11, and TFAP2D play a pivotal role in the development, identity, and PFC connectivity of these neurons. Using mouse models, we demonstrate that the absence of transcription factors SOX4 and SOX11 in post-mitotic ExNs dramatically reduces the size of the basolateral amygdala complex (BLC), claustrum, and PIR. SOX4 and SOX11 control BLC formation through direct regulation of *Tfap2d* expression. Cross-species analyses, including humans, identified conserved *Tfap2d* expression in developing ExNs of BLC, claustrum, paleocortex including PIR, and the associated transitional areas of the frontal, insular and temporal cortex. While the loss and haploinsufficiency of *Tfap2d* yield similar alterations in learned threat behaviors, differences emerge in the manifestation of *Tfap2d* dosage, particularly in terms of changes observed in BLC size and the connectivity pattern between the BLC and PFC. This underscores the significance of *Tfap2d* dosage in orchestrating developmental shifts in BLC-PFC connectivity and behavioral modifications reminiscent of symptoms of neuropsychiatric disorders. Together, these findings reveal key elements of a conserved gene regulatory network that shapes the development and function of crucial ventropallial ExNs and their PFC connectivity and offer insights into their evolution and alterations in neuropsychiatric disorders.

## Introduction

The claustro-amygdalar complex and the adjacent paleocortex play an indispensable role in integrating information crucial for emotional processing, memory consolidation, olfaction, and decision-making ^1-4^. This integration is made possible through the extensive long-range axonal connections between these ventrolateral pallial regions and various other brain regions ^2,3,5^. Notably, the reciprocal long-distance axonal connections of the glutamatergic excitatory neurons (ExNs) within the basolateral amygdala complex (BLC) and the piriform cortex (PIR; paleocortex) with the prefrontal cortex (PFC) are essential for the regulation of adaptive behavioral responses to environmental stimuli ^2-6^. Dysfunction in these connections has been closely associated with the pathophysiology of various neuropsychiatric disorders, many of which emerge early in life ^4,7-11^. During embryonic development, pallial-subpallial boundary (PSB) progenitors give rise to immature ExNs, which migrate via the lateral cortical stream (LCS) towards ventrolateral pallial structures like the PIR and specific components of the claustro-amygdalar complex, including the BLC ^12-21^. Despite previous important studies on the patterning and differentiation of these progenitors, the molecular mechanism governing the specification and connectivity of ExN progenies within these ventrolateral pallial structures has remained largely unexplored. In this study, we identify an evolutionary conserved gene regulatory network comprising of SOX4, SOX11, and TFAP2D, which orchestrates the post-mitotic, cell-autonomous specification of certain claustro-amygdalar and paleocortical ExN features, including their projections to the PFC, and their functional contributions.

## Results

### SOX4 and SOX11 Regulate Claustro-Amygdalar and Paleocortical Development

In prior observations, we noted that depletion of transcription factors (TFs), SOX4 and SOX11, from *Emx1*-expressing cortical progenitors resulted in significant developmental defects within the cerebral cortex ^22^. These defects included the loss of *Fezf2* expression, disruptions in the laminar specification of ExNs, and impaired formation of the corticospinal tract. Furthermore, it was observed that SOX4 and SOX11 are expressed in immature cerebral ExNs, yet their specific post-mitotic functions remain poorly understood beyond the aforementioned phenotypes ^22,24^. To delve deeper into these functions, we utilized *Neurd6-Cre*; *CAT-Gfp* mice and bred them with *Sox4* fl/fl and *Sox11* fl/fl mice. This breeding strategy allowed us to selectively eliminate the post-mitotic expression of both *Sox4* and *Sox11* from the cerebral ExNs (conditional double KO; cdKO) while simultaneously marking them with green fluorescent protein (GFP). Given the observed redundancy in the function of SOX4 and SOX11 ^22,23^, the *Sox4* cKO and *Sox11* cKO mice did not exhibit noticeable phenotypic defects compared to control littermates at postnatal day 0 (PD 0) (**Fig. 1a**, **Extended Data Fig. 1a-b, Extended Data Fig. 2a, 2c**). Similar to our previous observations ^22^, *Sox4; Sox11* cdKO brains showed phenotypic defects in corticospinal tract and cortical layer formation, suggesting that most of these effects were due to the post-mitotic functions of SOX4 and SOX11 (**Extended Data Fig. 1a-b**). Furthermore, in *Sox4*; *Sox11* cdKO mice, there was pronounced reduction observed in ventrolateral cortical structures, encompassing the BLC, claustrum and PIR (**Fig. 1a, Extended Data Fig. 1a**). Notably, these structures are known to express both *Sox4* and *Sox11* during development ^22^. Nissl staining, *in situ* hybridization, and immunostaining for markers outlining amygdala nuclei confirmed that *Sox4*; *Sox11* cdKO mice almost entirely lacked the BLC - comprising the basolateral (BLA), lateral (LA), and basomedial (BMA) nuclei - PD 0 (**Fig. 1a, Extended Data Fig. 2a**). Specifically, *Lmo3* and SATB1 comprehensively delineated the entire BLC; *Etv1* and *Mef2c* distinctly marked BLA and LA, respectively; and *Cyp26b1* prominently labeled both the BLA and the central amygdala nuclei ^18,22,25^. Additionally, NR4A2 (NURR1) immunostaining revealed reduction in the size of the endopiriform nucleus and claustrum in *Sox4*; *Sox11* cdKO brains (**Extended Data Fig. 2a**).

**Figure 1.**
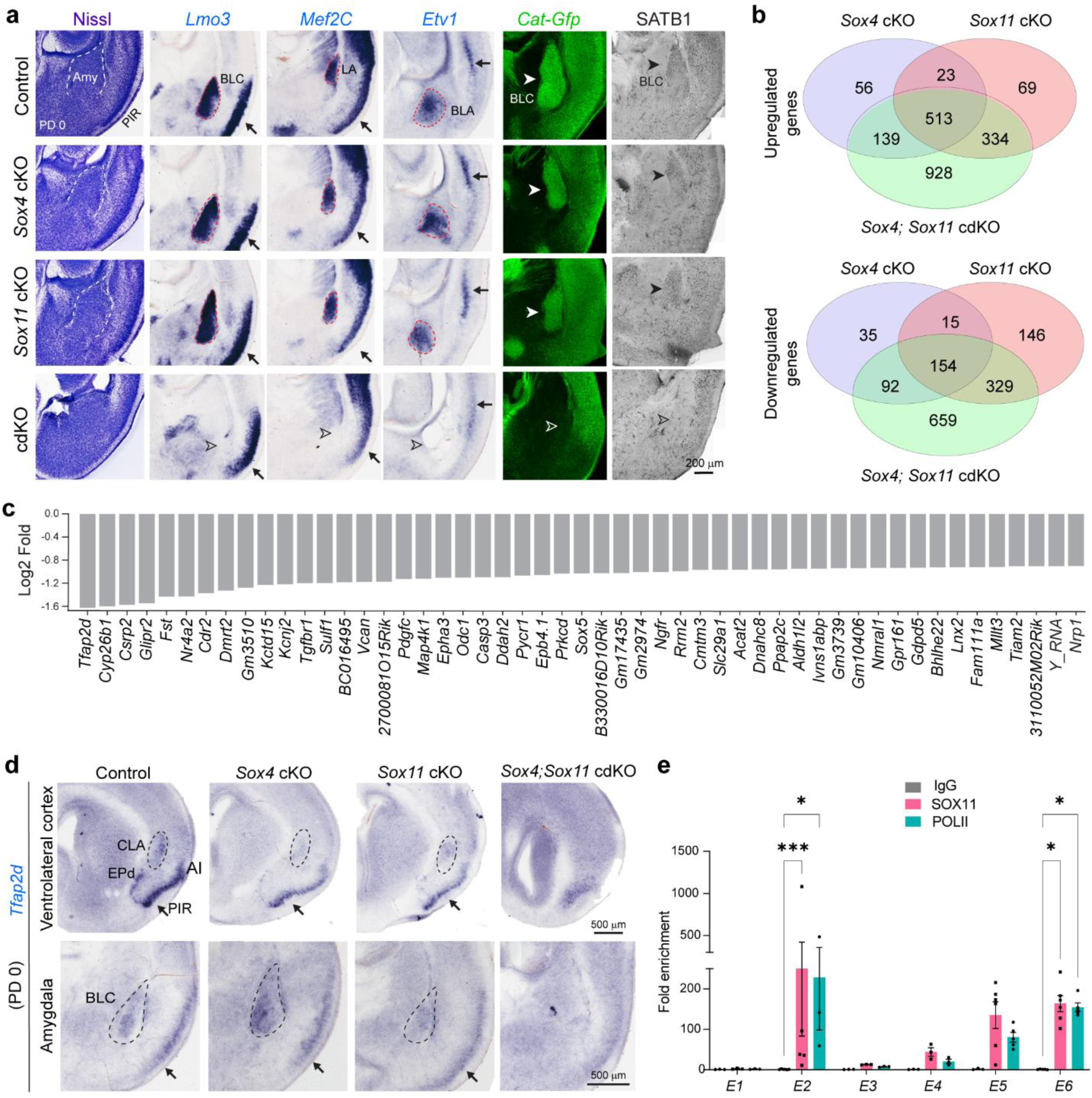
Regulation of Ventrolateral Pallial Development and *Tfap2d* Expression by SOX4 and SOX11. **a**) Coronal sections from the ventrolateral cortical region of controls (*Sox4* fl/+; *Sox11* fl/+; *Neurod6Cre*; *CAT-Gfp, or Sox4 fl/fl; Sox11fl/fl*), *Sox4* cKO (*Sox4* fl/fl; *Sox11* fl/+; *Neurod6Cre*; *CAT-Gfp*), *Sox11* cKO (*Sox4* fl/+; *Sox11* fl/fl; *Neurod6Cre*; *CAT-Gfp*), and *Sox4; Sox11* cdKO (*Sox4* fl/fl; *Sox11* fl/fl; *Neurod6Cre*; *CAT-Gfp*) mice brains at PD 0. The PIR and amygdala, specifically the BLC are highlighted by Nissl staining, in situ hybridization for *Lmo3*, *Mef2c,* and *Etv1*, show BLC, LA and BLA respectively and by immunostaining for GFP and SATB1 show BLC. Filled arrows and arrowheads point to normal phenotype and open arrowhead point to defective phenotypes in the knockouts. **b**) Venn diagram showing intersection of the number of differentially expressed genes in *Sox4* cKO, *Sox11* cKO and *Sox4; Sox11* cdKO brains as compared to the controls, as identified by PD 0 mouse cortex and amygdala using bulk-tissue RNA-seq (n= 3/genotype). We identified 3148, 1027, and 1583 differentially expressed genes between control samples and *Sox4; Sox11* cdKO, *Sox4* cKO, and *Sox11* cKO samples, respectively. **c**) Bar graph showing genes top 50 genes that have decreased in expression in *Sox4; Sox11* cdKO samples as compared to *Sox4* cKO, *Sox11* cKO and control samples. **d**) *In situ* hybridization of *Tfap2d* in the ventrolateral cortical region of the controls, *Sox4* cKO, *Sox11* cKO and *Sox4; Sox11* cdKO brains. In the control, *Sox4* cKO, and *Sox11* cKO brains expression of *Tfap2d* is seen in the CLA, AI, EPd, PIR, BLC. **e**) The quantitative PCR analysis of the chromatin selectively pulled down using antibodies for IgG (control), SOX11, or POLII, highlighting the enrichment against the input for six potential enhancers, labeled E1 through E6, located in and surrounding the *Tfap2d* gene locus. Mean ± s.e.m. depicted for n = 3/ antibody. Two-way ANOVA with multiple comparisons and Tukey’s correction was applied. p value <0.05*, p value <0.001*** Abbreviations: AI, Anterior Insula Amy, amygdala;; CLA, claustrum; EPd, endopiriform nucleus, dorsal part;.

### SOX4 and SOX11 Regulate Ventrolateral Pallial Expression of *Tfap2d*

To investigate downstream genes regulated by SOX4 and SOX11, we conducted bulk tissue RNA-seq analysis on pooled cerebral cortex and amygdala tissue obtained from PD 0 in *Sox4*; *Sox11* cdKO, *Sox4* cKO, *Sox11* cKO, and control littermates. We identified 958 and 659 genes that were uniquely upregulated or downregulated, respectively, in the *Sox4; Sox11* cdKO brains (**Fig. 1b**, **Extended Data Table 1-2**). Because several genes associated with neuroinflammation were among those upregulated in the *Sox4 Sox11* cdKO brains, we next assessed whether deletion of *Sox4*, *Sox11*, or both was associated with increased microglia and astroglia invasion (**Extended Data Table 1, Extended Data Fig. 2b**). Immunofluorescence with glial markers IBA1 and GFAP confirmed a significant increase in microglia and astroglia invasion, respectively (**Extended Data Fig. 2c**). In addition, TUNEL (terminal deoxynucleotidyl transferase biotin-dUTP) staining provided evidence for DNA fragmentation, suggesting cell damage (**Extended Data Fig. 2c**). These data corroborate previously reported functions of SOX4 and SOX11 in neuron differentiation and survival ^23,24^.

Because the severe defects in the ventrolateral pallium were seen only in the *Sox4; Sox11* cdKO, we focused on the 659 genes that were exclusively downregulated in these mice (**Fig. 1b, Extended Data Table 2**). Many of the top downregulated genes have been previously shown ^25-29^, to be enriched in claustro-amygdalar cortex or paleocortical structures, including *Cyp26b1*, *Glipr2*, *Fst* and *Nr4a2* (**Fig. 1c, Extended Data Fig. 2a**). The topmost downregulated gene in *Sox4; Sox11* cdKO mice encodes the transcription factor, TFAP2D (also known as AP-2 Delta) (**Fig. 1c, Supplementary Table 2**), which we have previously shown to be enriched in the human and non-human primate amygdala ^26-29^. While TFAP2D has previously been implicated in the development of the dorsal midbrain ^30^ and retina ^31^, its role in the development of the amygdala or other ventral pallidal structures has not been studied. *In situ* hybridization and analysis of transcriptomic datasets revealed that the mouse *Tfap2d* exhibits elevated expression levels in developing ExNs in the BLC and the neighboring ventrolateral pallial structures, such as the PIR, anterior olfactory nucleus, endopiriform nucleus, anterior insula, and claustrum (**Fig. 1d**, **Fig. 2e and Extended Data Fig. 5c, Extended Data Table 2**). In accordance with the results of the RNA-seq analysis, the expression of *Tfap2d* was nearly absent in these structures of *Sox4*; *Sox11* cdKO mice when assessed at PD 0 (**Fig. 1d**).

**Figure 2.**
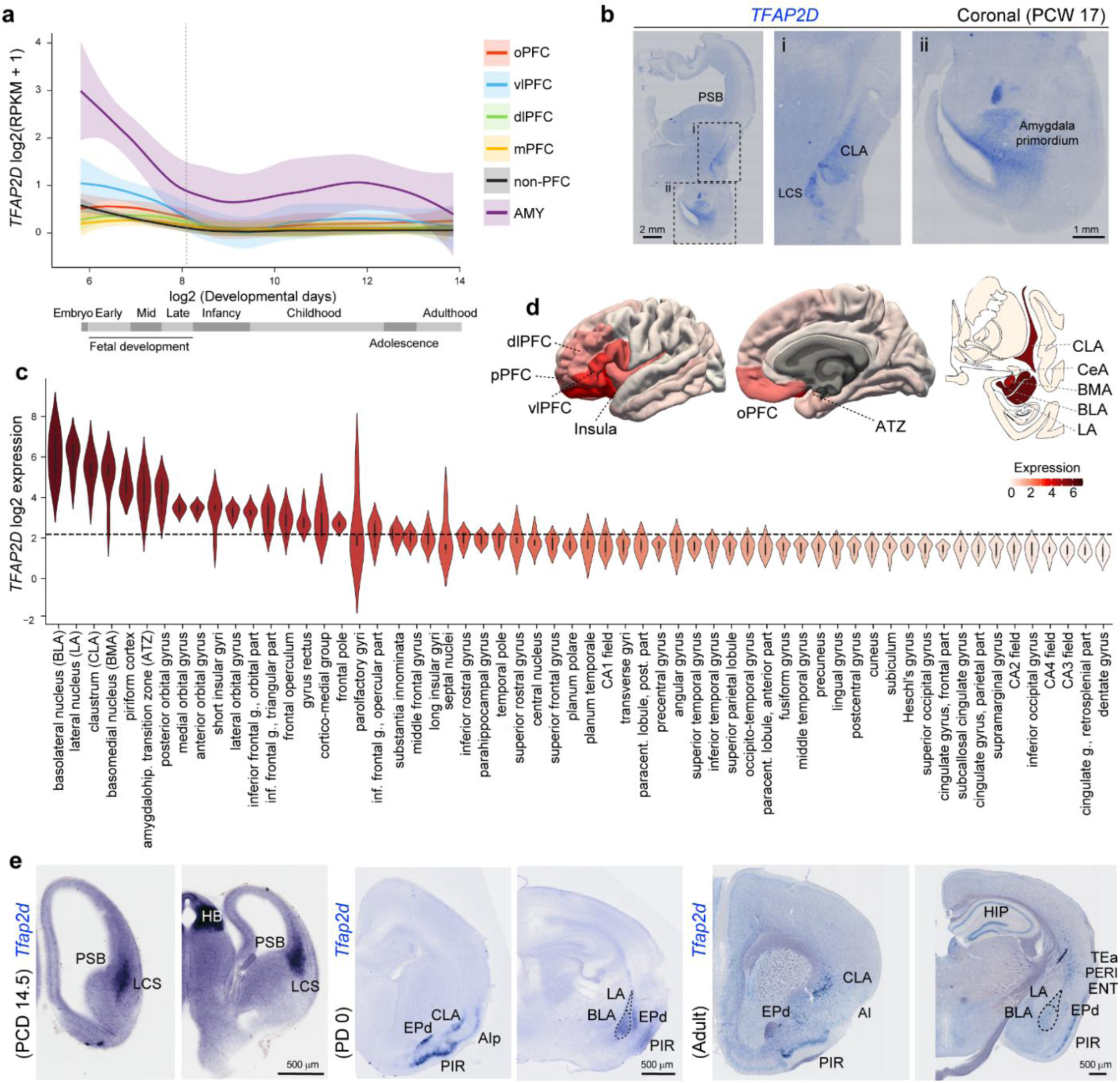
Conserved Expression Pattern of *TFAP2D* in Human and Mouse Brain. **a)** Plot depicting the expression of *TFAP2D* in across developmental ages in the human amygdala (AMY), and Orbital (o), ventral (v), dorsolateral (dl), medial (m) and non-PFC (prefrontal cortex) ^26,27^. **b)** Coronal section of a human post-conception week (PCW 17) brain showing the expression of *TFAP2D* within the LCS, CLA and amygdala primordia. **c)** *TFAP2D* expression pattern across frontal cortical regions, ventrolateral cortical structures, and subcortical regions in adult human brain as derived from publicly available microarray data ^34^. The dotted line represents the mean of the *TFAP2D* expression across all the regions represented here. **d)** Human brain images showing *TFAP2D* expression gradients across adult brain areas. To visualize the *TFAP2D*, correspondence between the Desikan-Killiany atlas and the Allen Brain Atlas probes was established. Subcortical regions were visualized using data from http://atlas.brain-map.org. **e)** *Tfap2d* in situ hybridization in mouse coronal sections at PCD 14.5, PD 0 and adult wildtype mice. Abbreviations: AIp, AI, posterior part; ATZ, amygdalohippocampal transition zone; ENTI, entorhinal cortex; HB, hebanula; PERI, perirhinal cortex; pPFC, polar PFC; TEA, temporal association area.

To explore whether SOX4 and SOX11 directly regulate the expression of *Tfap2d*, we conducted an integrated analysis, cross-referencing chromatin immunoprecipitation (ChIP)-seq data on H3K27ac ^32^, a marker of active enhancers and promoters, and accessible chromatin data from ATAC-seq ^33^ in the mouse forebrain spanning post-conception day (PCD) 11.5 to PD 0 with predicted SOX binding motifs and SOX11 ChIP-seq data ^24^ (**Extended Data Fig. 3a**). Through this analysis, we identified six putative enhancers associated with *Tfap2d* expression that also contain SOX11 binding motifs, which also showed increased accessibility starting at PCD 13.5, which coincides with the migration of immature amygdalar ExNs ^16,18^. To validate these *in silico* findings, we conducted ChIP followed by real-time PCR (RT-PCR) using a SOX11 antibody that we validated in *Sox11* cKO brains (**Extended Data Fig. 3b**). We failed to identify a suitable anti-SOX4 antibody for immunohistochemistry or ChIP assays. Our ChIP results revealed strong binding of SOX11 to two of the six identified putative enhancers, which we designated as E2 and E6 (**Fig. 1e**). Remarkably, these regulatory regions also exhibited robust binding of POLII, indicating their active involvement in *Tfap2d* transcription. Consistent with these findings, *Sox4*, *Sox11* and *Tfap2d* are co-expressed by developing ExNs within the BLC (**Extended Data Fig. S4 c-d, Extended Data Fig. S6 a-d**). Collectively, these findings indicate that at least SOX11 directly regulates the expression of *Tfap2d* in the ventral pallium.

### Evolutionary Conserved Expression of *Tfap2d* in Claustro-Amygdalar and Paleocortical ExNs

Our previous transcriptomic analyses ^26-29^ revealed an enrichment of *TFAP2D* in the human and non-human primate prenatal amygdala that undergoes a steady decline until infancy and persists at lower levels thereafter (**Fig. 2a**). We corroborated *TFAP2D* expression in the human brain at post-conception week 17 (PCW 17), specifically within the LCS, as well as within the claustro-amygdalar primordia, employing *in situ* hybridization (**Fig. 2b**). Furthermore, our analysis of microarray datasets from the midfetal and adult human brains ^34,35^ revealed a gradient of *TFAP2D* expression, with higher levels observed in BLA, LA, claustrum, BMA, and PIR, followed by the peripaleocortical subdivisions of the frontal, insular and temporal cortex, and the orbito-lateral transitional neocortical PFC areas (**Fig. 2c-d, Extended Data Fig. 4a**). We further confirmed the expression pattern of *TFAP2D* in the amygdala and claustrum by ddPCR (**Extended Data Fig. 4b**). To identify which human amygdala cell types express *TFAP2D*, we used publicly available human amygdala single nuclear RNA sequencing (snRNA-seq) ^36^ data which demonstrated that *TFAP2D* is only highly expressed within the class of amygdala ExNs that also co-expressed *SOX4* and *SOX11* as well as other classic ExN markers such as *TBR1*, *NEUROD2*, and *SLC17A6* (**Extended Data Fig. 4c-d**).

Similarly to humans, our examination of the RNA-seq dataset sourced from the developing and adult macaque and chimpanzee brain regions ^28,29^ uncovered pronounced prenatal expression of *TFAP2D* in the amygdala, which progressively diminishes with advancing age (**Extended Data Fig. 5a**). To ascertain the conservation of *TFAP2D* expression patterns in the developing ventrolateral pallial structures across species, we conducted *in situ* hybridization experiments in both rhesus macaques and mice. Our investigation revealed *TFAP2D* expression in the macaque LCS and within the amygdala primordium at post conception day (PCD) 105 (**Extended Data Fig. 5b**). In mouse, we performed *in situ* hybridization at PCD 14.5, PD 0, and PD 60. At PCD 14.5, we observed *Tfap2d* within the neurons migrating along LCS, and in the amygdala primordium (**Fig. 2f, Extended Data Fig. 5c**). Notably, *Tfap2d* expression was not observed in the proliferative zones at the PSB in either mouse or macaque (**Fig. 2e, Extended Data Fig. 5b-c**). This finding indicates that *Tfap2d* starts to be expressed within immature post-mitotic neurons that undergo migration along the LCS. At both PD 0 and PD 60, *Tfap2d* is expressed within the BLC, claustrum, and anterior insula (**Fig. 2e, Extended Data Fig. 5c**), like in humans (**Fig. 2c**), as well as within the endopiriform nucleus and PIR (**Fig. 2e, Extended Data Fig. 5c**). At PD 60 we also observed a sporadic expression pattern of *Tfap2d* within the deep layers of the anterior insula, insular gustatory, temporal association, and auditory cortex (**Extended Data Fig. 5c)**. We further corroborated this expression pattern in mouse using publicly available cerebrum snRNA-seq datasets ^37^ (**Extended Data Fig. 5d-e**). Using RNAscope, we detected the expression of *Tfap2d* within the nidopallium and mesopallium of the chicken forebrain on embryonic day (E) 17, including within the PIR and the arcopallial amygdala (**Extended Data Figure 5f**). These regions are homologues of the mammalian ventrolateral pallial structures, including the BLC, claustrum, and PIR. This evidence, indicates that *Tfap2d* expression is highly conserved across a broad spectrum of species, including humans, non-human primates, mice, and chickens.

To evaluate the cell type-specific expression of *TFAP2D* across various species, we conducted a reanalysis of publicly available amygdala snRNA-seq datasets in human, macaque, mouse, and chicken ^38^. Our findings reveal heightened and consistent expression of *Tfap2d* in the BLC and major ExN subclasses across all species profiled (**Extended Data Fig. 6a-d**). Within human, *TFAP2D* has the highest expression in the *LAMP5* ExN clusters (**Extended Data Fig. 6a**), which were dominant in the human BLC ^38^. Additionally, we observed the expression of *SOX4*, *SOX11* and *TFAP2D* in the paralaminar nucleus (PL) in humans (**Extended Data Fig. 6a**), which has been previously shown to harbor immature SOX11-positive ExNs in the adulthood ^39^. In mice, the *Rspo* ExN subclass had the highest expression of *Tfap2d* and *Sox11* (**Extended Data Fig. 6c**), a cell type known to regulate anxiety-like behaviors in mice ^40^. The comparable *TFAP2D* expression pattern across human, macaque, mouse, and chicken in the ExNs within ventrolateral pallial structures suggests that there may be a conserved TFAP2D-dependent regulatory network for the specification of ExNs populating these regions.

### *Tfap2d* Expression in ExNs is Necessary for BLA and LA Formation

To examine the impact of *Tfap2d* deletion on the development of ventrolateral pallial structures, we analyzed mice with a targeted disruption of *Tfap2d*. Initially, we investigated *Tfap2d* lacZ mice, where a β-galactosidase-neomycin cassette, interrupting exon 1 of the *Tfap2d* gene, induces a complete truncation of the protein and abolishes *Tfap2d* expression ^30^ (**Extended Data Fig. 7a**). These mice were bred to generate *Tfap2d* knockout (KO), heterozygous (Het), and wildtype (WT) control groups. Given the previously reported expression and function of *Tfap2d* in the superior colliculus ^30^, we also developed *Tfap2d* floxed mice by inserting a floxed cassette within introns 1 and 3 of the *Tfap2d* gene, creating a conditional allele (*Tfap2d* fl) (**Extended Data Fig. 8a-b, Extended Data Table 3**). For targeted deletion of *Tfap2d* in post-mitotic cerebral ExNs, we crossed these mice with *Neurod6Cre*; *CAT-Gfp* mice, yielding conditional knockout groups (*Tfap2d* cKO, *Tfap2d* cHet, and WT control) that express a GFP reporter specifically in cerebral ExNs. In *Tfap2d* KO and cKO mice, *Tfap2d* expression was absent in ventrolateral pallial structures, including the BLC, PIR, claustrum and endopiriform nucleus. Expression in the superior colliculus remained unaffected in cKO mice (**Fig. 3a, Extended Data Fig. 8c**). Animals of all genotypes including the Hets and KOs survived to adulthood.

**Figure 3.**
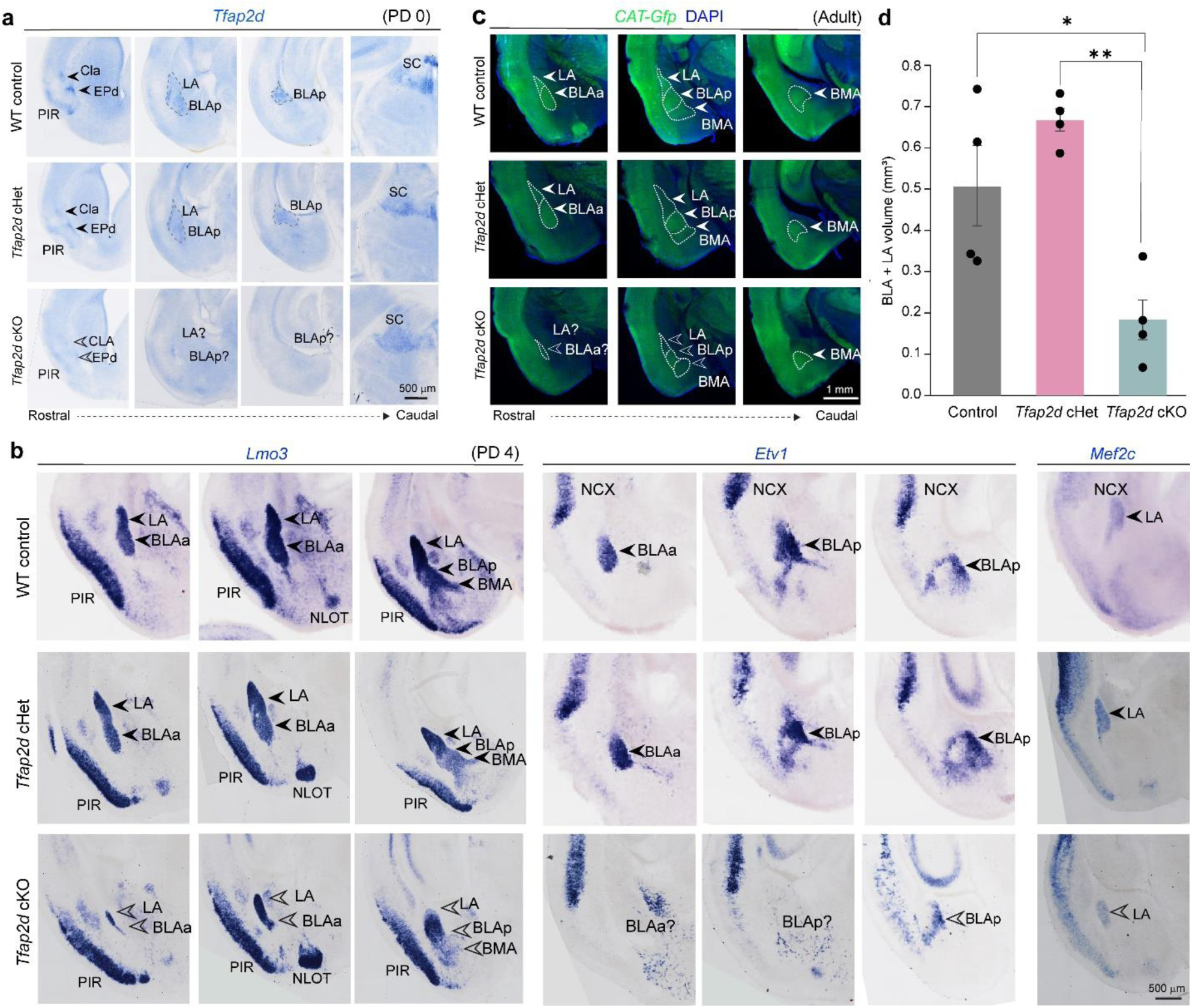
Post-mitotic Deletion of *Tfap2d* in Cerebral ExNs Leads to Reduced BLC. **a**) Coronal sections of WT control (*Tfap2d* +/+; *Neurod6Cre*; *Cag-CAT-Gfp* or *Tfap2d fl/fl*) *Tfap2d* cHet (*Tfap2d* fl/+; *Neurod6Cre*; *Cag-CAT-Gfp*) and *Tfap2d* cKO (*Tfap2d* fl/fl; *Neurod6Cre*; *Cag-CAT-Gfp*) carrying deletion of *Tfap2d* in cerebral post-motitic ExN. Depicting the specificity of the deletion *Tfap2d* expression is reduced in the ventrolateral cortical structures-CLA, EPd, PIR, BLC but not in the Sc in the midbrain. Arrowheads point to normal expression pattern and open arrowhead point to loss of *Tfap2d* expression in the *Tfap2d* cKO. **b**) Coronal sections of the ventrolateral cortical regions of the WT Control, *Tfap2d* cHet and *Tfap2d* cKO brains showing the expression of *Lmo3*, *Etv1* and *Mef2c* by *in situ* hybridizations at PD 4. Reduction in the size of BLC is seen by LMO3, the most affected region is the *Etv1* labelled BLA. **c**) Coronal sections of WT Control, *Tfap2d* cHet and *Tfap2d* cKO showing the cytoarchitecture of the brain in adults. *Gfp* labeling depicts reduced BLC size, mostly due to BLA and LA, in the *Tfap2d* cKO as compared to the WT control, and *Tfap2d* cHet mice **d**) Stereological measurements of the volume of BLA + LA in the adult WT control, *Tfap2d* cHET and *Tfap2d* cKO brains. Mean ± s.e.m. depicted. Two-way ANOVA with Tukey’s correction was applied. ** and * represent p values < 0.01, <0.05 respectively (n= 4/genotype). BLAa, BLA, anterior part; BLAp, BLA, posterior part; NLOT, nucleus of lateral olfactory tract; NCX, neocortex; SC, superior colliculus.

To assess whether ventrolateral pallial development is altered following *Tfap2d* loss, we conducted a comprehensive analyses of perinatal mouse brains. No significant defects were observed in neocortical or archicortical structures, including their GFP-labeled long-range subcerebral and interhemispheric projections in the *Tfap2d* cKO mice (**Extended Data Fig. 9a**), aligning with the restricted expression pattern of *Tfap2d* to claustro-amygdalar and paleocortical ExNs. However, Nissl and AChE staining, as well as GFP expression, indicated a dramatic reduction in the size of BLC in *Tfap2d* KO and cKO mice, respectively (**Figure 3c-d, Extended Data Fig. 7e**).

Further, immunostaining for TBR1 and SATB1 (**Extended Data Fig. 9b**) and *in situ* hybridization for *Lmo3*, *Etv1*, and *Mef2c* (**Figure 3b, Extended Data Fig. 7b**), key markers of amygdalar ExNs, confirmed a near-complete loss of the BLA, partial loss in the LA, but no obvious loss in the BMA in both *Tfap2d* KO and cKO mice compared to their respective Hets and WT controls. Immunostaining for NR4A2, a marker of claustral ExNs, showed no noticeable differences in claustrum between *Tfap2d* KO, cKO, and WT control mice (**Extended Data Fig. 7b, Extended Data Fig. 9c**). In adults, no appreciable anatomical abnormalities were observed in other ventrolateral pallial structures in either *Tfap2d* cKO or KO mice (**Fig. 3c, Extended Data Fig. 7c-d**). These findings indicate that *Tfap2d* plays an essential cell-autonomous role in the post-mitotic development of LA and BLA ExNs.

### Disruption of BLC-PFC Connectivity by *Tfap2d* Loss or Haploinsufficiency

The BLC ExNs possess extensive long-range connections to multiple brain regions, including the PFC, thalamus, and hippocampus ^41^. These connections are important for the BLC to encode salient cues from the environment and transmit the relevant signals to downstream structures ^2-6^. Prompted by the observed reduction in the BLC in *Tfap2d*-deficient mice, we embarked on an investigation to delineate the influence of *Tfap2d* on BLC’s long-range neural projections. Utilizing Diffusion Tensor Imaging (DTI) on controls, *Tfap2d* Het, and KO brains, we discovered a marked impairment in BLC-PFC connectivity in *Tfap2d* KO mice (**Fig. 4a**). Intriguingly, despite a significant reduction in BLC-PFC connectivity in *Tfap2d* Het brains compared to WT controls, these brains exhibited significantly increased connectivity relative to *Tfap2d* KO brains, indicating a dosage-dependent, intermediate impact of *Tfap2d* haploinsufficiency on BLC-PFC connections (**Fig. 4c**). No appreciable differences were observed in the BLC-thalamus or BLC-hippocampus connections across the genotypes (**Fig. 4a-c**). Further analyses to ascertain whether *Tfap2d* loss uniformly affects neocortical connections, we specifically examined contralateral cortical-cortical connections in the somatosensory cortex, primary motor cortex, and PFC (**Fig. 4a-c**), expectedly revealed no significant discrepancies across genotypes, suggesting a targeted effect of *Tfap2d* deficiency on specific BLC-related neural pathways.

**Figure 4.**
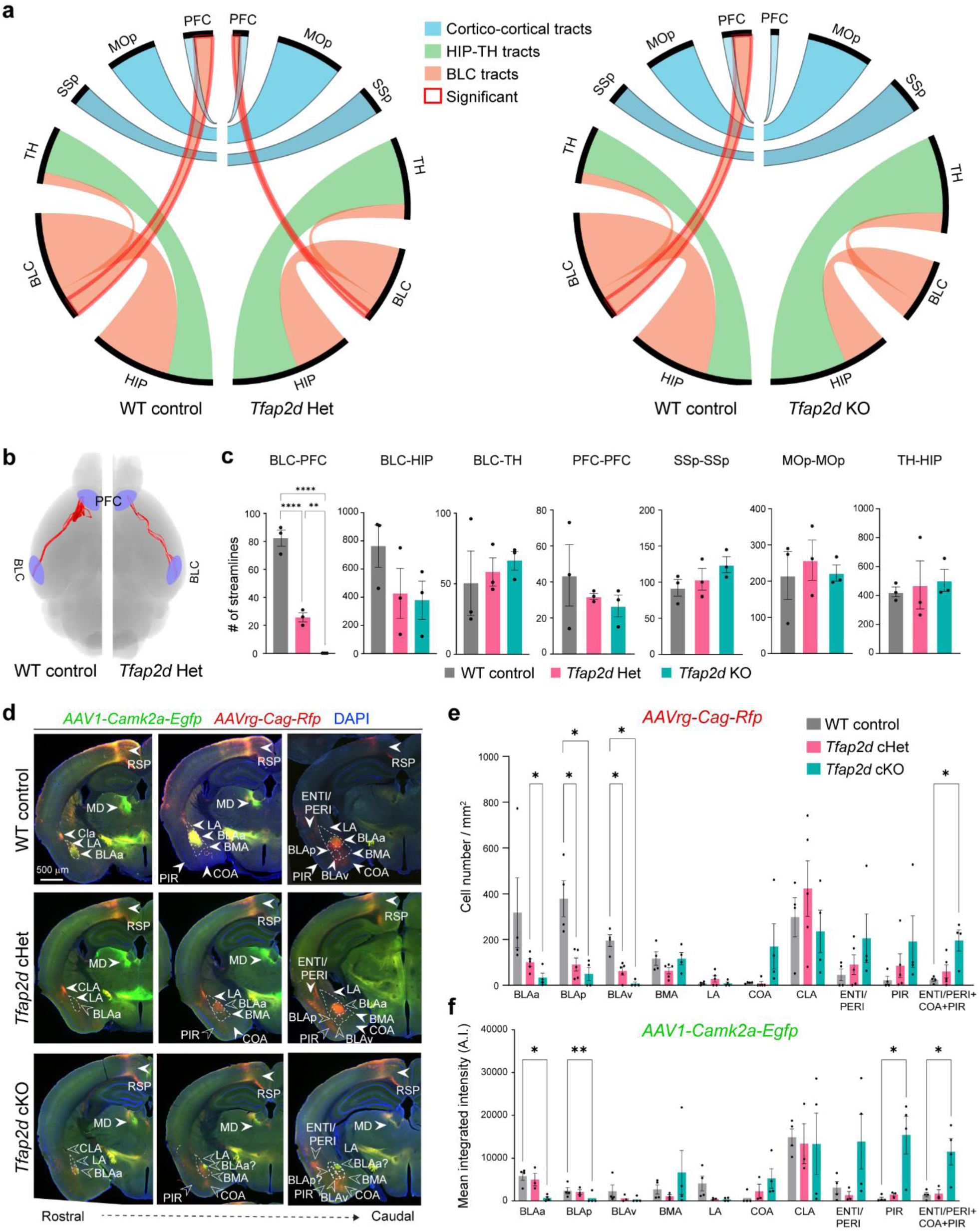
*Tfap2d* Loss or Haploinsufficiency Alters Connections between BLC and PFC. **a)** Representation of the number of streamlines (NOS) generated as a connectivity measurement between the BLC and mPFC, BLC and HIP, BLC and TH, at PD 120-180 of the *Tfap2d* KO animals using Diffusion Tensor Imaging (DTI). In *Tfap2d* Het and *Tfap2d* KO brains, there is a significant (red outline) decrease in NOS generated between the BLC and mPFC as compared to WT control. The NOS between other regions are not significantly different between genotypes (n = 3 per genotype). **b)** Visualization of the streams connecting the BLC and mPFC in the WT control and *Tfap2d* Het. **c**) Quantification of the NOS between BLC and mPFC, BLC and HIP, BLC and TH and between contralateral cortical areas in WT control, *Tfap2d* Het and *Tfap2d* KO mice. Paired t-test: ****, ** < 0.0001, <0.01, respectively; n = 3 per genotype. **d**) Representative images of the WT control, *Tfap2d* cHet and *Tfap2d* cKO brains whose efferent and afferent projections from the mPFC were traced using *AAV1-Camk2a-Egfp* (anterograde, green) and *AAVrg-Cag-Rfp* (retrograde, red) at PD 90. Open arrows indicate reduced or misdirected projections from and to the mPFC. **e**) Quantifications of the retrograde tracings showing number of cells projecting to mPFC from the BLAa, BLAp, BLAv, COA, PIR and ENTI/PERI in the *Tfap2d* cKO and *Tfap2d* cHet. Mean ± s.e.m. depicted. Two-way ANOVA with Tukey’s correction was applied. ** and * represent p values, < 0.01, <0.05 respectively (n = 4 (WT), 5 (cHet) and 4 (cKO). **f**) Quantifications of the anterograde tracings show reduced intensity of projections from PFC to the BLAa, BLAp, BLAv and increased intensity to COA, PIR and ENTI/PERI areas in the *Tfap2d* cKO and *Tfap2d* cHet mice, as compared to WT control mice. Mean ± s.e.m. depicted. Two-way ANOVA with Tukey’s correction was applied. ** and * represent p values < 0.01, <0.05 respectively (n= 4 (WT), 3 (cHet) and 4 (cKO). BLAv, BLA, ventral part; MOp, primary motor cortex; MD, mediodorsal nucleus of the thalamus; RSP, retrosplenial cortex; SSp, primary somatosensory cortex; TH, Thalamus.

To further corroborate the deficits in BLC-PFC connectivity observed by DTI, we utilized adeno-associated virus (AAV)-based tract tracing in *Tfap2d*-deficient mice. Given the significant reduction in size of the BLC in *Tfap2d*-deficient animals, we introduced both anterograde and retrograde AAV vectors (*AAV1-Camk2a-Egfp* and *AAVrg-Cag-Rfp*, respectively) into the medial PFC (mPFC), broadly targeting both the prelimbic (PL) and infralimbic (ILA) areas. The mPFC was targeted because it showed significant connectivity alterations in the DTI analysis. In WT control animals, the expected projection patterns between the BLA and mPFC were observed (**Fig. 4d-f, Extended Data Fig. 10a-c**). Contrastingly, in both *Tfap2d* cKO and KO mice, a notable reduction in the number of neurons projecting from the BLA to the PFC and reduction in the intensity of projections from PFC to BLA was evident (**Fig. 4d-f, Extended Data Fig. 10a-c**). In addition to the reduced BLA-mPFC connections, a dispersed projection pattern from the cortical amygdalar area (COA), PIR, and perirhinal cortex to the PFC was observed in *Tfap2d* KO and cKO mice compared to WT counterparts (**Fig. 4d-f, Extended Data Fig. 10a-c**). This shift in projection patterns suggests a compensatory redistribution of BLA-PFC projections to other ventrolateral cortical regions in *Tfap2d*-deficient animals. Consistent with the DTI observations, *Tfap2d* Het and cHet mice also exhibited significant deviations in the BLA-mPFC projection pattern relative to WT mice (**Fig. 4d-f, Extended Data Fig. 10a-c**). These deviations varied among individuals, with some displaying abnormal PIR-PFC and COA-PFC projections and others showing a general decrease in BLA-PFC connectivity. Collectively, these findings underscore that the absence of one or both *Tfap2d* alleles results in altered BLA-mPFC connections, highlighting the critical role of *Tfap2d* gene dosage in maintaining proper neural circuitry.

### *Tfap2d*-Induced BLC Deficits Compromise Threat-Response

The BLC plays a crucial role in encoding the valence of environmental cues, transmitting this critical information to other brain regions tasked with triggering behavioral responses ^6^. To investigate the impact of *Tfap2d*-driven deficits in BLC development and adult exploratory behavior, we conducted a battery of behavioral tests on adult *Tfap2d* KO and cKO animals, comparing their responses to those of their respective WT and Het counterparts (**Extended Data Figs. 11a-b and 12a-f**). Among these tests, the open field and zero maze assessments were employed to evaluate whether disruptions in BLC formation attributed to *Tfap2d* loss are linked with exploratory behaviors. The findings reveal that both *Tfap2d* KO and cKO subjects exhibit a tendency to spend less time in the center and more time along the periphery of the open field, relative to their WT and Het counterparts, indicating a potential association between *Tfap2d*-dependent BLC development deficits and altered defensive responding (**Extended Data Fig. 11a, 12a-c**). Distance traveled was the same for all genotypes, indicating that there were no gross motor deficits in either the *Tfap2d* KO or *Tfap2d* cKO animals (**Extended Data Fig. 11a, 12c**). In the zero-maze test, *Tfap2d* KO, and *Tfap2d* cKO animals spent more time in the closed arm of the zero maze as compared their respective WT groups (**Extended Data Fig. 11b, 12d-f**). *Tfap2d* cHet mice also spent more time in the closed arm as compared to WT control (**Extended Data Fig. 11b**). *Tfap2d* KO animals exhibited fewer entries to the light side of the light-dark box test than either WT or Het animals but displayed no differences in the forced swim test or tail suspension test, tests related to depression-like behaviors (**Extended Data Fig. 11c-e**). These data demonstrate that *Tfap2d*-dependent deficits in BLC formation and connectivity leads to alterations in threat behaviors in adults, in which exploratory behaviors are diminished.

The BLC is recognized for its pivotal role in facilitating the learning of associations between neutral environmental cues and adverse outcomes, such as fear conditioning ^2-6^. To test the effects of *Tfap2d*-driven BLC-related deficits on negative association learning, we performed fear conditioning (**Fig. 5a-c, Extended Data Fig. 13a-c**). Both *Tfap2d* KO and *Tfap2d* cKO animals learned this association at the same rate as their respective Het and WT groups (**Fig. 5a, Extended Data Fig. 13a**). During the contextual conditioning test, where animals were placed in the same context as in training but without sound presentation, *Tfap2d* Het animals spent significantly less time freezing as compared to the WT animals (**Extended Data Fig. 13b**). Similarly, *Tfap2d* cHet animals also spent significantly less time freezing than WT control and *Tfap2d* cKO animals (**Fig. 5b**). During the cued conditioning test, *Tfap2d* KO and *Tfap2d* cKO animals, as well as their respective Hets, displayed consistently higher levels of freezing as compared to WT animals, indicating that loss or haploinsufficiency of *Tfap2d* causes alterations in the expression of learned threat responses (**Fig. 5c, Extended Data Fig. 13c**). Moreover, the behavioral patterns seen in the *Tfap2d* mice align with observed effects of haploinsufficiency in other AP2 family TFs on learned behaviors ^42^, although the functions of other AP2 genes on BLC development are not known. Taken together, these results suggest that haploinsufficiency or loss of *Tfap2d* leads to nuanced changes associated with the interpretation of salient cues that may stem from the alterations in the BLC-related circuitry.

**Figure 5.**
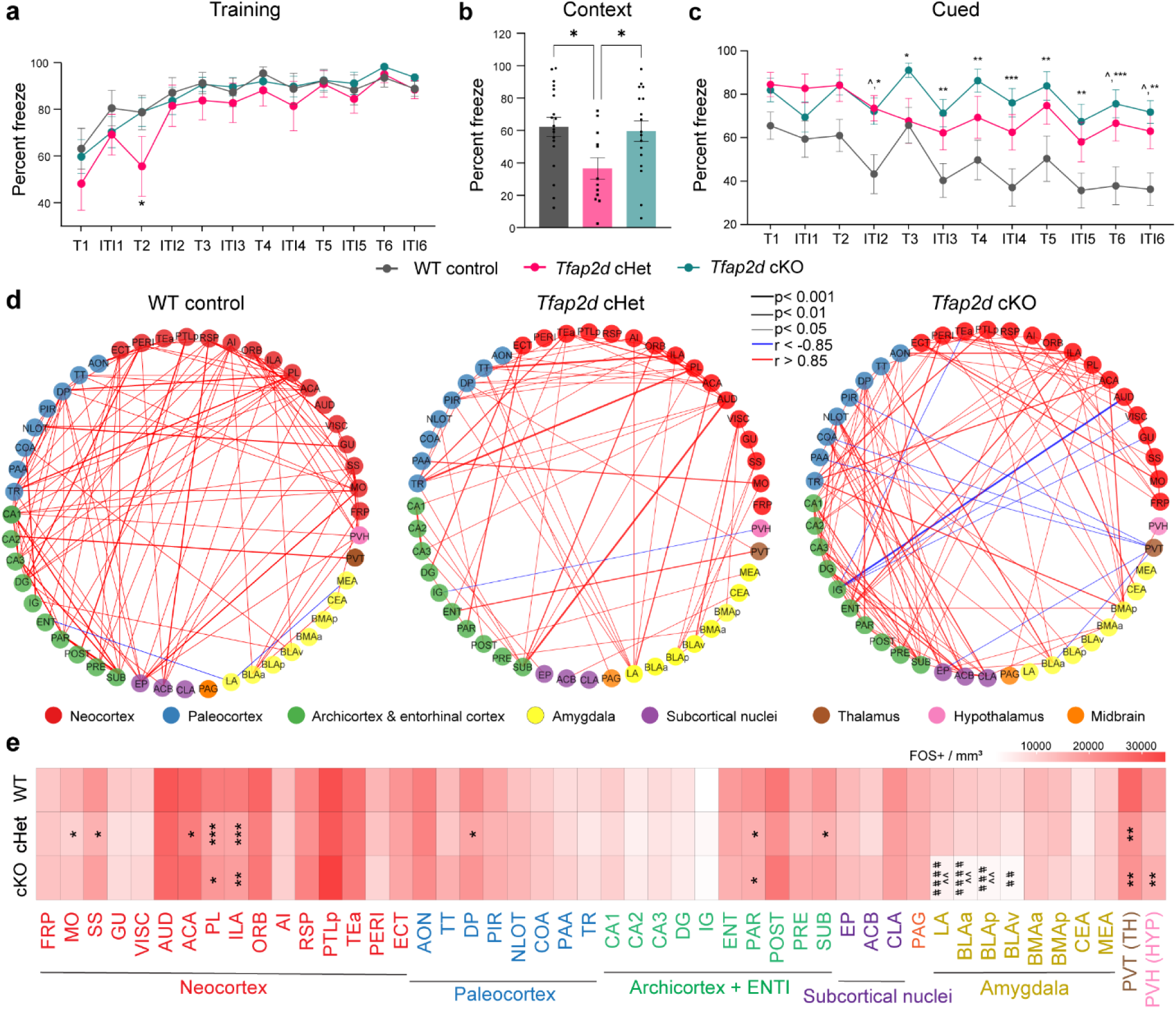
*Tfap2d* Loss or Haploinsufficiency Increases Threat Responding in Learned Behaviors and Alters Functional Connectivity. **a)** Line plot showing the learning response during the fear conditioning test of the WT control, *Tfap2d* cHet and *Tfap2d* cKO mice as the percentage of time (y-axis) animals freeze during and inter tone interval (ITI) and tone (75 decibels, 20 seconds) (T) followed by a 2 sec shock at 0.5mA (x-axis). **b**) Bar plot showing conditioned responses of the WT control, *Tfap2d* cHet and *Tfap2d* cKO mice as percentage of time the animals froze (y-axis) while in the training context. One-way ANOVA with Tukey’s correction was applied. ** and * represent p values < 0.01, <0.05 respectively. **c**) Line plot showing cued responses of the WT control, *Tfap2d* cHet and *Tfap2d* cKO mice as percentage of time (y-axis) animals freeze during T and ITI without shock (x-axis). Mean ± s.e.m. depicted. Two-way ANOVA with repeated measures and multiple comparison Tukey’s correction was applied. ***, ** and * represent p values <0.001, < 0.01, <0.05 respectively between WT and cKO, whereas ^ represents p value < 0.05 between WT and cHet. n = 16 (WT control), 12 (Tfap*2d* cHet) and 16 (*Tfap2d* cKO). **d**) Network graph depicting the interregional correlation between brain regions as measured by the number of FOS positive cells. in the WT control, *Tfap2d* cHet and *Tfap2d* cKO. The brain regions are represented as nodes, and their significant correlation (p value <0.01 and r > 0.85) scores as edges. The color of the node depicts the broader brain area. The color of the edges indicates the direction of correlation (red and blue indicate positive and negative correlation, respectively). **e**) Heatmap depicting FOS positive cells per mm^3 detected within the different brain regions of the WT control, *Tfap2d* cHet and *Tfap2d* cKO brains isolated after 60-90 min of cued test. Mean ± s.e.m. depicted. Two-way ANOVA with Tukey’s correction was applied. ***, **, * represents p values < 0.001, <0.01, <0.05 respectively, for (n= 4/ genotype). For LA, BLAa, BLAp, BLAv, ^# # # #^p, ^# # #^p, ^# #^p represents p values <0.0001, < 0.001, <0.01, respectively for WT vs cKO and ^^^^p for p values <0.01 or cHet vs cKO. Abbreviations: Refer to Extended Data Table 4 for 5d-e.

During the cued conditioning test, behaviors are determined by synchronized communication among multiple brain regions ^43^. To discern the patterns of coordinated brain activity associated with fear expression and that may be impaired by *Tfap2d* haploinsufficiency or loss, we computed interregional correlations of FOS density, a measure of gross neuronal activity. We correlated FOS density between regions that are associated with cued fear such as the amygdala, neocortex, hippocampus, and thalamus (**Fig. 5d, Extended Data Fig. 14a**).

We noticed that significant correlations between cortical regions and the hippocampus formation in the WT were reduced or altered in both *Tfap2d* cHet and *Tfap2d* cKO. In WT control animals the FOS densities of LA and PL were negatively correlated, whereas this trend reverses in the *Tfap2d* cHet animals. However, in the *Tfap2d* cKO mice, we didn’t observe any significant correlation between LA and PL (**Fig. 5d, Extended Data Fig. 14a**). Importantly, there is a shift within the amygdala in the FOS correlation matrix in the *Tfap2d* cKO animals compared to WT animals. In WT animals, BLA FOS density is correlated with IL; however, in *Tfap2d* cKO animals, which are missing the BLA, we observe a significant correlation between the perirhinal cortex, entorhinal cortex, and BMA with PL, ILA and ACA. These results corroborate with the circuit alterations seen in the T*fap2d* cKO mice. However, these altered circuits do not seem to compensate for the loss of the BLA, as evidenced by the altered functional connectivity and disrupted threat responses observed in the cued tests. These data indicate that *Tfap2d* haploinsufficiency or loss led to changes in FOS density correlations between threat-associated regions. Notably, there exist distinct differences in amygdala and PFC FOS density correlations between WT, *Tfap2d* cHet and *Tfap2d* cKO mice. To highlight significant correlations in FOS density we generated network graphs in which nodes represent brain regions, and edges represent the significant correlations that surpassed a set threshold (**Fig. 5d**). The network analysis indicated a substantial reduction in the significant correlations among brain regions that are involved in threat responding in the *Tfap2d* cHet while in the *Tfap2d* cKO we observed a shift in this connectivity compared to the WT (**Fig. 5d**). These differences are consistent with the observed circuit deficits and indicate that *Tfap2d* haploinsufficiency alters functional connectivity in a different manner than a total loss of *Tfap2d*.

Concurrently, we observed significantly fewer FOS positive cells in the ILA and PL mPFC in *Tfap2d* KO and Het mice, as well as *Tfap2d* cKO and cHet mice, as compared to their respective WT groups (**Fig. 5e, Extended Data Fig. S13 d-e**). We note that this FOS pattern was not observed within the amygdala where there were fewer FOS positive cells in LA, BLAa, and BLAp in *Tfap2d* KO and *Tfap2d* cKO as compared to both their respective Het and WT mice. These data suggests that even early-life haploinsufficiency of *Tfap2d*, though not sufficient to impair BLC formation, may be enough to lead to differences in the expression of, but not the acquisition of, learned defensive behaviors which are also reflected in differences in FOS levels across the brain.

## Discussion

This study addresses a critical question at the juncture of neuroscience and evolutionary biology, exploring the molecular mechanisms that underpin the development and evolution of the ventrolateral pallium. Our analysis yields three principal findings. First, we present evidence demonstrating the indispensable role of the SOX4 and SOX11-dependent post-mitotic gene regulatory network in the proper development of ventrolateral pallial ExNs, particularly within the BLC complex. This network orchestrates the expression of genes with highly specific patterns within the developing claustro-amygdalar complex and paleocortex, which are crucial for neuronal development and connectivity. These insights build upon and extend the known functions of these TFs in neuronal specification ^24^ including broadly neocortical and archicortical ExNs ^22^. Here, we highlight a specific regulatory node essential for the molecular specification of ventrolateral pallial ExN development.

Second, our findings identify TFAP2D as a pivotal TF within this ventrolateral pallial regulatory node. Its expression, highly specific to ventrolateral pallial ExNs and contingent upon SOX4 and SOX11 activity, including direct SOX11 binding at enhancers in the TFAP2D locus, underscores its critical role. This study significantly expands our understanding of TFAP2D’s role beyond its previously documented expression in the human and non-human primate amygdala ^26-29^, revealing its broader pattern across paleocortical and the associated mesocortex. The nuanced role of TFAP2D is further evidenced by the delicate balance of its gene dosage, which profoundly affects the development of these structures and results in altered behaviors and connectivity patterns, particularly between the BLC and PFC.

Thirdly, we demonstrate that these regulatory networks and gene expression patterns are conserved across multiple species, suggesting an evolutionary preserved molecular mechanism. Interestingly, the presence of an Alu cassette in the human *TFAP2D* locus, absent in other primates ^44^, hints at potential human-specific variations in *TFAP2D* expression, a factor that could play a role in shaping our unique characteristics within these brain regions.

These findings also emphasize the nuanced influence of *Tfap2d* gene dosage on developmental trajectories, BLC-PFC connectivity, and behaviors that resemble alterations observed in certain neurodevelopmental and neuropsychiatric disorders. Accordingly, mutations in SOX4 and SOX11 cause intellectual disability and other neurodevelopmental alterations ^45,46^, and variants within the TFAP2D locus are associated with bipolar disorder and emotional dysregulation ^47-50^, opening new avenues for investigating the molecular mechanisms underlying BLC and paleocortical development in the context of disease. This exploration may uncover how alterations in these processes contribute to neural traits and disease conditions, offering novel insights into their pathogenesis.

## Acknowledgements

We would like to acknowledge and thank Veronique Lefebvre for providing *Sox4* flox and *Sox11* flox mice. We thank Alvaro Duque and Yale Macaque Brain Resource (MBRC) for use of their Aperio CS2 scanner and stereology system. Funding for the MBRC is provided by NIH (grant RO1MH113257 to Alvaro Duque). We would also like to acknowledge the Yale Rodent Behavior Analysis Facility supported by the Kavli Institute for Neuroscience for providing assistant with the behavioral tests. R.K. was supported by 1F32NS117780-01 during this research period. G.S. is supported by grants MS20/00064 (ISCIII-MICINN/FEDER), PID2019-104700GA-I00 and PID2022-140137NB-I00 (/AEI/10.13039/501100011033), Fundació LaMarató de TV3 and NIH grant HG010898 (to G.S and N.S.). X.D.M. is supported by the fellowship PRE2020-093064 funded by MCIN/ AEI/10.13039/501100011033. This work was supported by NIH grants and MH077681 (to Y.S.M. and M.R.P.), NS095654, MH106934, MH116488, MH110926, and MH129981 and the Simons Foundation (to N.S.).

## Contributions

N.K., F.O.G., M.P., J.S., and N.S. conceived the study and methodology. N.K., R.K., F.O.G., M.P., J.S., and N.S. designed the research. N.K. primarily conducted the experimental work, with N.K. and R.K. overseeing data quality control and analyses. M.M. generated the whole-body *Tfap2d* KO mice, while F.O.G. conducted the preliminary analysis of these mice. N.K. generated the *Tfap2d* floxed mice and designed and conducted RNA-seq experiments. N.K., R.K., F.O.G., D.F., M.Y., I.C., D.E., and M.B. contributed to mouse breeding and genotyping. J.M., Y.L., H.C. X.D.M, G.S., D.A., N.K., and R.K. performed the computational analyses. N.K., T.S.K., F.O.G, and J.S. carried out behavioral tests. K.P., Y.S.M., and M.R.P. provided guidance on behavioral tests and analysis. H.H. conducted DTI imaging and provided insights into data interpretation. D.A., T.Z. and N.Sa. analyzed DTI data. A.S., and M.P. performed post-mortem human, chicken, and macaque tissue gene expression analyses. N.K., R.K. and N.S. wrote the manuscript, including the creation of figures and tables, with input from all authors during result discussion, implications assessment, and manuscript review.

