## Supplementary Figures for "Molecular Specification of Claustro-Amygdalar and Paleocortical Neurons and Connectivity"

### Extended Data Figures

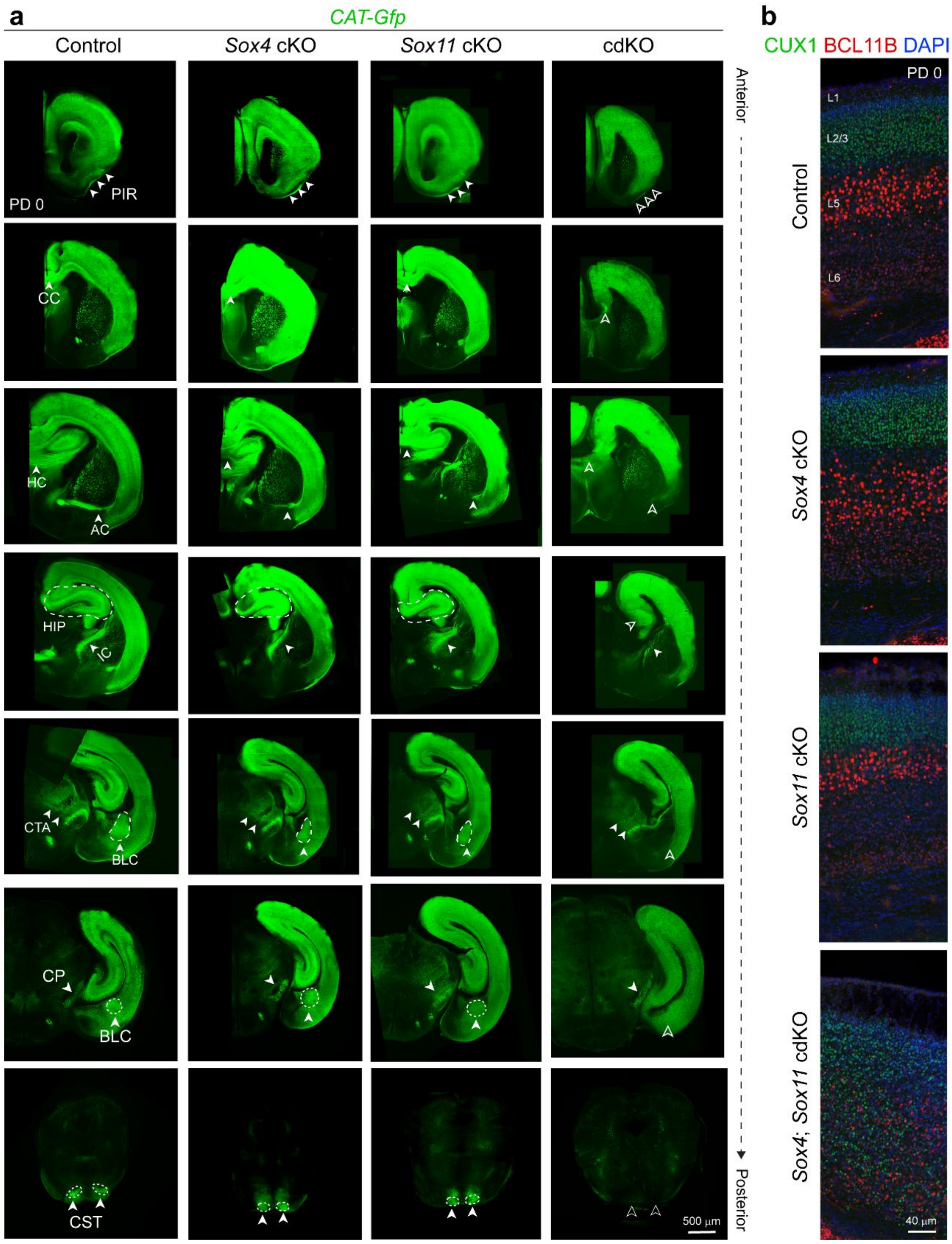

**Extended Data Figure 1. *Sox4* and *Sox11* Loss Disrupts ExN Projections and Laminar Organization**

**a**) Serial sections of control, *Sox4* cKO, *Sox11* cKO and *Sox4; Sox11* cdKO mice brains at PD 0. Filled arrows point to normal phenotype and open arrowhead point to defective phenotypes in the knockouts. **b**) Representative images illustrate immunostaining for CUX1 and BCL11B in the cerebral cortex, highlighting disrupted laminar organization and decreased density nuclei with pronounced BCL11B immunosignal in the *Sox4*; *Sox11* cdKO brains, in contrast to control, Sox4 cKO, and *Sox11* cKO samples. AC, anterior commissure; CC, corpus callosum; CP, cerebral peduncles; CST, corticospinal tract; CTA, corticothalamic axons; HC, hippocampal commissure; HIP, hippocampus; IC, internal capsule; L, layer.

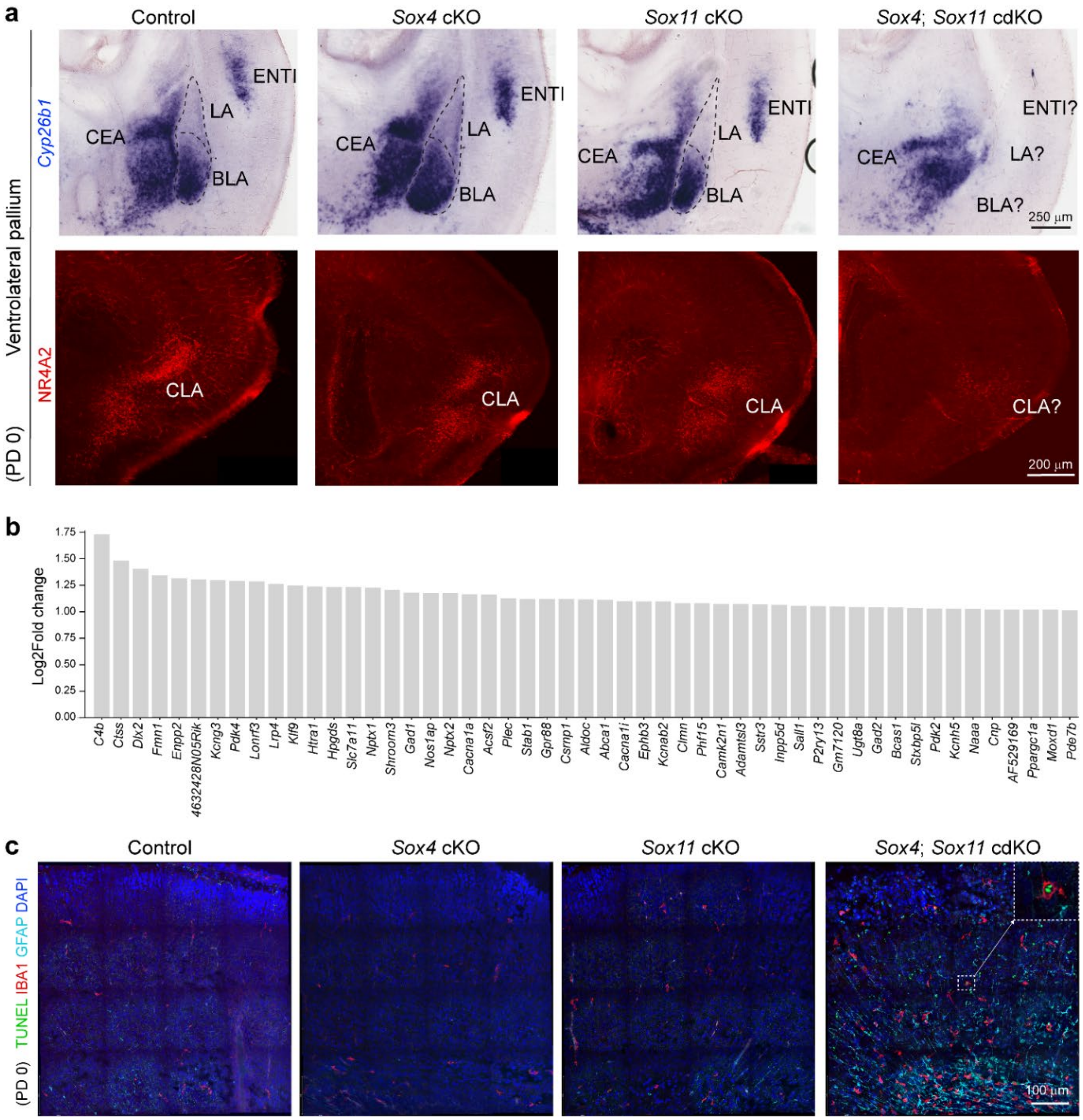


**Extended Data Figure 2. *Sox4* and *Sox11* Deletion Impairs Cell Survival, Affecting the Development of Ventrolateral Pallial Structures**

**a**) Coronal sections of the *in situ* hybridization for *Cyp26b1* (upper panel) and immunostaining for NR4A2 validating their downregulation in the ventrolateral cortical structures- including BLC, CLA, and ENTI in *Sox4*; *Sox11* cdKO as compared to control, *Sox4* cKO, and *Sox11* cKO brains. In all genotypes, *Cyp26b1* expression is robust in the CEA, the GABAergic inhibitory center of amygdala. **b**) Bar graph showing top 50 genes that have increased in expression in *Sox4; Sox11* cdKO samples as compared to *Sox4* cKO, *Sox11* cKO and control within cortex and amygdala at PD 0. **c**) Representative images showing the immunostaining for GFAP and IBA1 along with TUNEL assay to depict the cell damage, astrocytes and microglia invasion in the *Sox4; Sox11* cdKO as compared to the control and *Sox4* cKO, *Sox11* cKO cortices at PD 0. CEA, central nucleus of the amygdala; CLA, claustrum; ENTI, entorhinal cortex; LA, lateral nucleus of the amygdala.

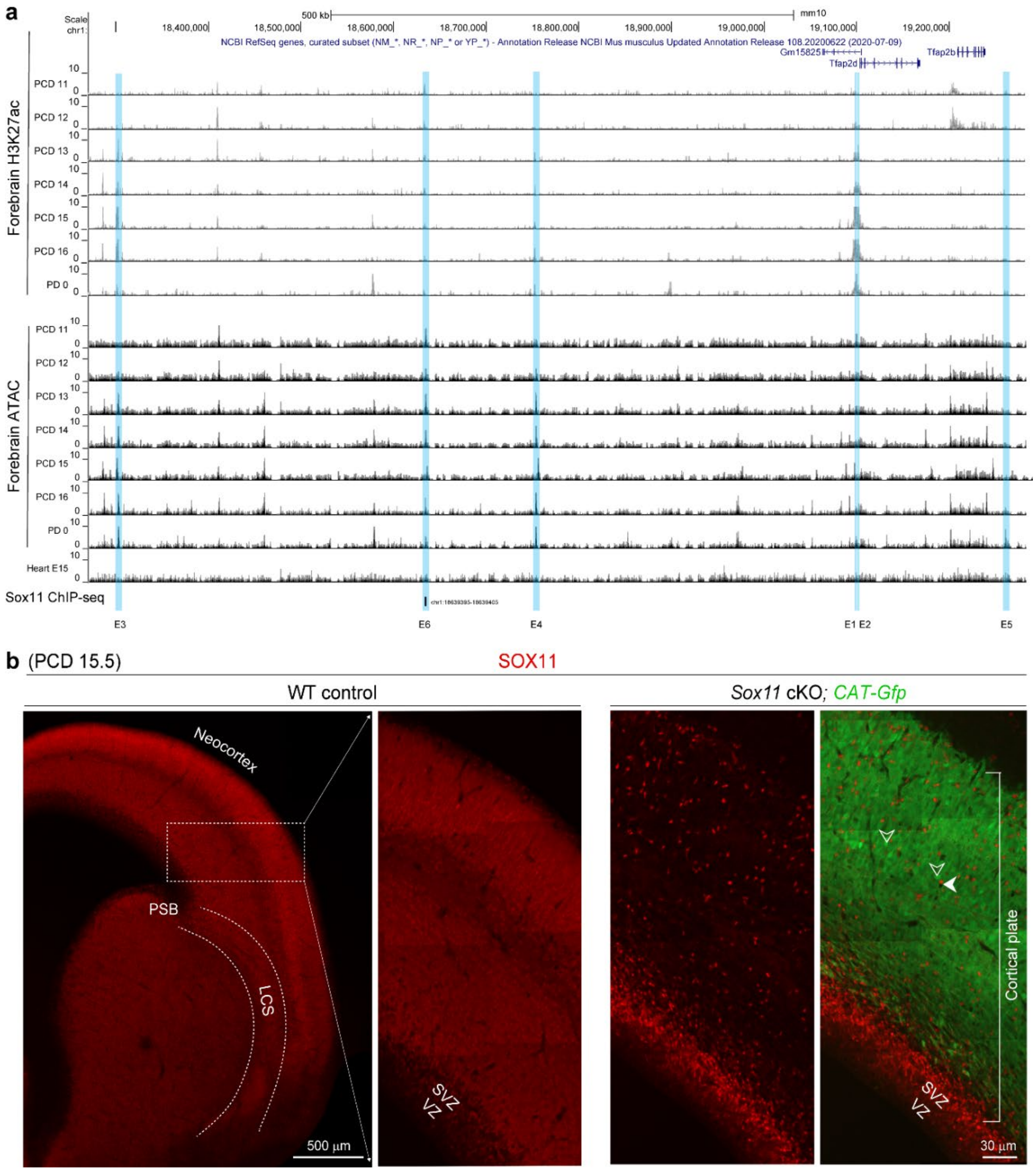


**Extended Data Figure 3. SOX11 Regulate *Tfap2d* Expression by Targeting its Enhancer Region**

**a**) Line graphs showing the forebrain H3K27ac[32] and ATAC peaks[33] and SOX11 ChIP-seq sites[24] from ages PCD 11-16, and PD 0 at the *Tfap2d* genomic locus ATAC peaks from heart (bottom tract) samples depict that the mechanism is forebrain specific. The blue boxes highlight putative enhancers (*E*) harboring predicted binding sites for SOX11, spanning from *E1* to *E6*, with ChIP-seq confirmation of binding specifically within *E6*. **b**) Validation of the anti-SOX11 antibody involved examining coronal sections of PDC 15.5 brains. In WT control brains, SOX11 immunostaining (red) was observed, while in the SOX11 cKO, expression from all GFP+ ExNs was lost (indicated by open arrowheads). Filled arrowheads point to SOX11 immunopositive nuclei within the cortical plate and in SVZ. ATAC; Assay for Transposase-Accessible Chromatin. SVZ, subventricular zone; VZ, ventricular zone; WT, wildtype.

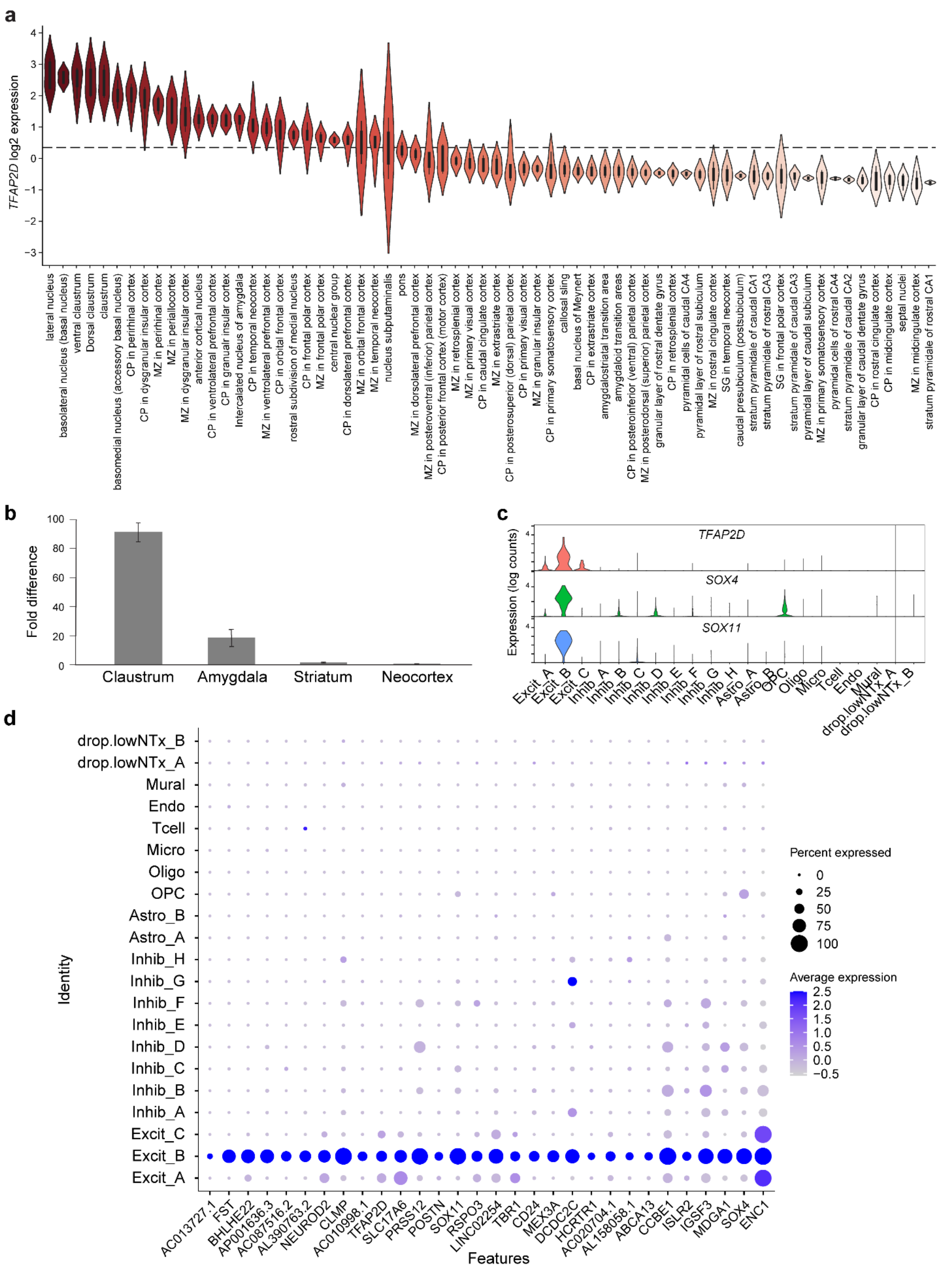
a
TFAP2D log2 expression
lateral nucleus
basolateral nucleus (basal nucleus)
ventral claustrum
Dorsal claustrum
claustrum
basomedial nucleus (accessory basal nucleus)
CP in perirhinal cortex
CP in dysgranular insular cortex
MZ in perirhinal cortex
MZ in periallocortex
MZ in dysgranular insular cortex
anterior cortical nucleus
CP in ventrolateral prefrontal cortex
CP in granular insular cortex
Intercalated nucleus of amygdala
CP in temporal neocortex
MZ in ventrolateral prefrontal cortex
CP in orbital frontal cortex
rostral subdivision of medial nucleus
CP in frontal polar cortex
MZ in frontal polar cortex
central nuclear group
CP in dorsolateral prefrontal cortex
MZ in orbital frontal cortex
MZ in temporal neocortex
nucleus subputaminalis
pons
MZ in dorsolateral prefrontal cortex
MZ in posteroventral (inferior) parietal cortex
CP in posterior frontal cortex (motor cortex)
MZ in retrosplenial cortex
MZ in primary visual cortex
CP in caudal cingulate cortex
MZ in extrastriate cortex
CP in posterosuperior (dorsal) parietal cortex
CP in primary visual cortex
MZ in granular insular cortex
CP in primary somatosensory cortex
callosal sling
basal nucleus of Meynert
CP in extrastriate cortex
amygdalostriatal transition area
amygdaloid transition areas
CP in posteroinferior (ventral) parietal cortex
MZ in posterodorsal (superior) parietal cortex
granular layer of rostral dentate gyrus
CP in retrosplenial cortex
pyramidal cells of caudal CA4
pyramidal layer of rostral subiculum
MZ in rostral cingulate cortex
SG in temporal neocortex
caudal presubiculum (postsubiculum)
stratum pyramidale of caudal CA1
stratum pyramidale of rostral CA3
SG in frontal polar cortex
stratum pyramidale of caudal CA3
pyramidal layer of caudal subiculum
MZ in primary somatosensory cortex
pyramidal cells of rostral CA4
stratum pyramidale of caudal CA2
granular layer of caudal dentate gyrus
CP in rostral cingulate cortex
CP in midcingulate cortex
septal nuclei
MZ in midcingulate cortex
stratum pyramidale of rostral CA1
b
Fold difference
Claustrum
Amygdala
Striatum
Neocortex
c
Expression (log counts)
TFAP2D
SOX4
SOX11
Excit_A
Excit_B
Excit_C
Inhib_A
Inhib_B
Inhib_C
Inhib_D
Inhib_E
Inhib_F
Inhib_G
Inhib_H
Astro_A
Astro_B
OPC
Oligo
Micro
Tcell
Endo
Mural
drop.lowNTx_A
drop.lowNTx_B
d
Identity
drop.lowNTx_B
drop.lowNTx_A
Mural
Endo
Tcell
Micro
Oligo
OPC
Astro_B
Astro_A
Inhib_H
Inhib_G
Inhib_F
Inhib_E
Inhib_D
Inhib_C
Inhib_B
Inhib_A
Excit_C
Excit_B
Excit_A
AC013727.1
FST
BHLHE22
AP001636.3
AC087516.2
AL390763.2
NEUROD2
CLMP
AC010998.1
TFAP2D
SLC17A6
PRSS12
POSTN
SOX11
RSPO3
LINC02254
TBR1
CD24
MEX3A
DCDC2C
HCRTR1
AC020704.1
AL158058.1
ABCA13
CCBE1
ISLR2
IGSF3
MDGA1
SOX4
ENC1
Features
Percent expressed
0
25
50
75
100
Average expression
2.5
2.0
1.5
1.0
0.5
0.0
−0.5

**Extended Data Figure 4. Expression of *TFAP2D* in the Human Brain**

**a**) *TFAP2D* expression pattern across frontal cortical regions, ventrolateral cortical structures, and subcortical regions in prenatal human brain[35]. The dotted line represents the mean of *TFAP2D* expression across all the regions represented here. X-axis depicts the regions and y-axis depicts the log 2 value for the expression pattern. **b**) Barplot showing the expression of *TFAP2D* in the claustrum, amygdala, striatum and neocortex detected by dd-PCR in adult human tissue (n = 3). **c**) Violin plot generated using publicly snRNA-seq available data[36] showing *TFAP2D*, *SOX4,* and *SOX11* expression in the ExNs (Excit) of the human amygdala with highest enrichment in the Excit_B class. y-axis depict the $log_2$ expression values and x-axis depicts the different classes of neurons within the human amygdala. **d**) Dot plot showing the percentage of cells that express the top 30 genes within the Excit_B class. Color represents the average expression level and size of the circle represent the percentage of cells.

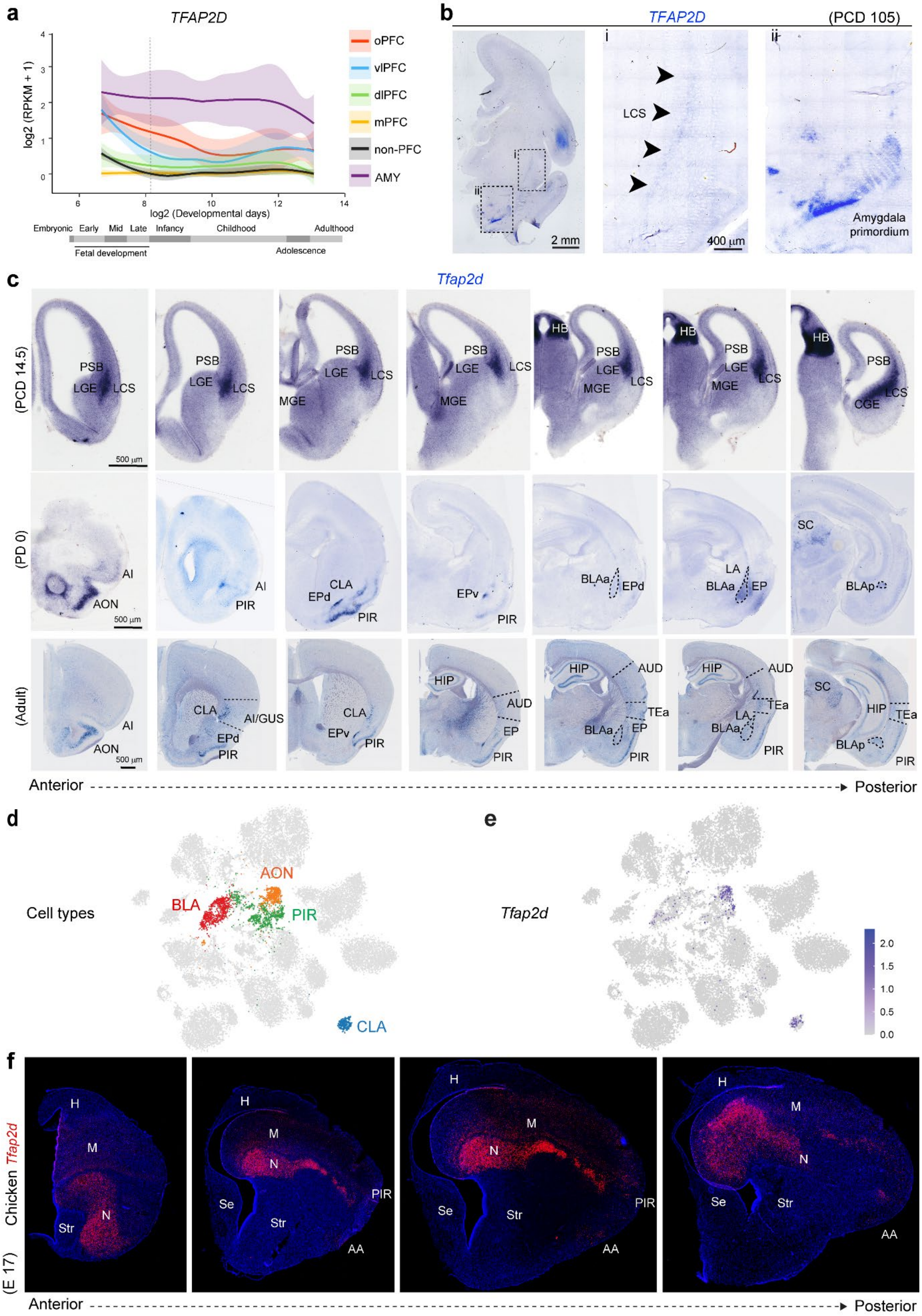
a
TFAP2D
log2 (RPKM + 1)
log2 (Developmental days)
oPFC
vlPFC
dlPFC
mPFC
non-PFC
AMY
Embryonic
Early
Mid
Late
Infancy
Childhood
Adulthood
Fetal development
Adolescence
b
TFAP2D
(PCD 105)
i
ii
LCS
2 mm
400 µm
Amygdala primordium
c
Tfap2d
(PCD 14.5)
(PD 0)
(Adult)
PSB
LGE
LCS
MGE
CGE
HB
500 µm
AI
AON
PIR
CLA
EPd
EPv
BLAa
LA
EP
SC
BLAp
AI/GUS
HIP
AUD
TEa
Anterior
Posterior
d
Cell types
BLA
AON
PIR
CLA
e
Tfap2d
2.0
1.5
1.0
0.5
0.0
f
Chicken Tfap2d
(E 17)
H
M
N
Str
Se
PIR
AA
Anterior
Posterior

**Extended Data Figure 5. Expression of *Tfap2d* in the Macaque and Mouse Brain**

**a**) Plot of *TFAP2D* expression across developmental ages in the macaque AMY, OFC, DFC, VFC, MFC and nonPFC areas within the forebrain [28,29]. **b**) Coronal section of a macaque brain at PCD 105 showing *TFAP2D* expression in the LCS, and amygdala primordium detected by *in situ* hybridization. **c**) Representative serial coronal sections showing *Tfap2d* expression within wildtype mouse PCD 14.5, PD 0 and adult brains detected by *in situ* hybridization. At PCD 14.5, *Tfap2d* expression is restricted to the LCS and Hb primordium. At PD 0 and in adults *Tfap2d* expression is seen in the BLC, CLA, EPd, EPv PIR, and SC. In adults, deep layer neurons in the AUD, GUS, and TEA show sporadic expression of *Tfap2d*. Representative images from this panel are used in Fig. 2e. **d**-**e**) UMAP showing the subclasses of glutamatergic neurons in the adult mouse cerebrum (left) and the expression of *Tfap2d* within the subclasses representing the BLC, PIR, AON, CLA [37]. f) Coronal sections of the chicken brain showing *Tfap2d* (red) expression in the nidopallium and arcopallial amygdala detected by RNAscope. AA, arcopallium amygdala; AON, anterior olfactory nucleus; AUD, auditory cortex; EPv, EP, ventral part; GUS, gustatory cortex; H, hyperpallium; LGE, lateral ganglionic eminence; M, mesopallium; MGE, medial ganglionic eminence; N, nidopallium; SC, superior colliculus; Se, septum; Str, striatum.

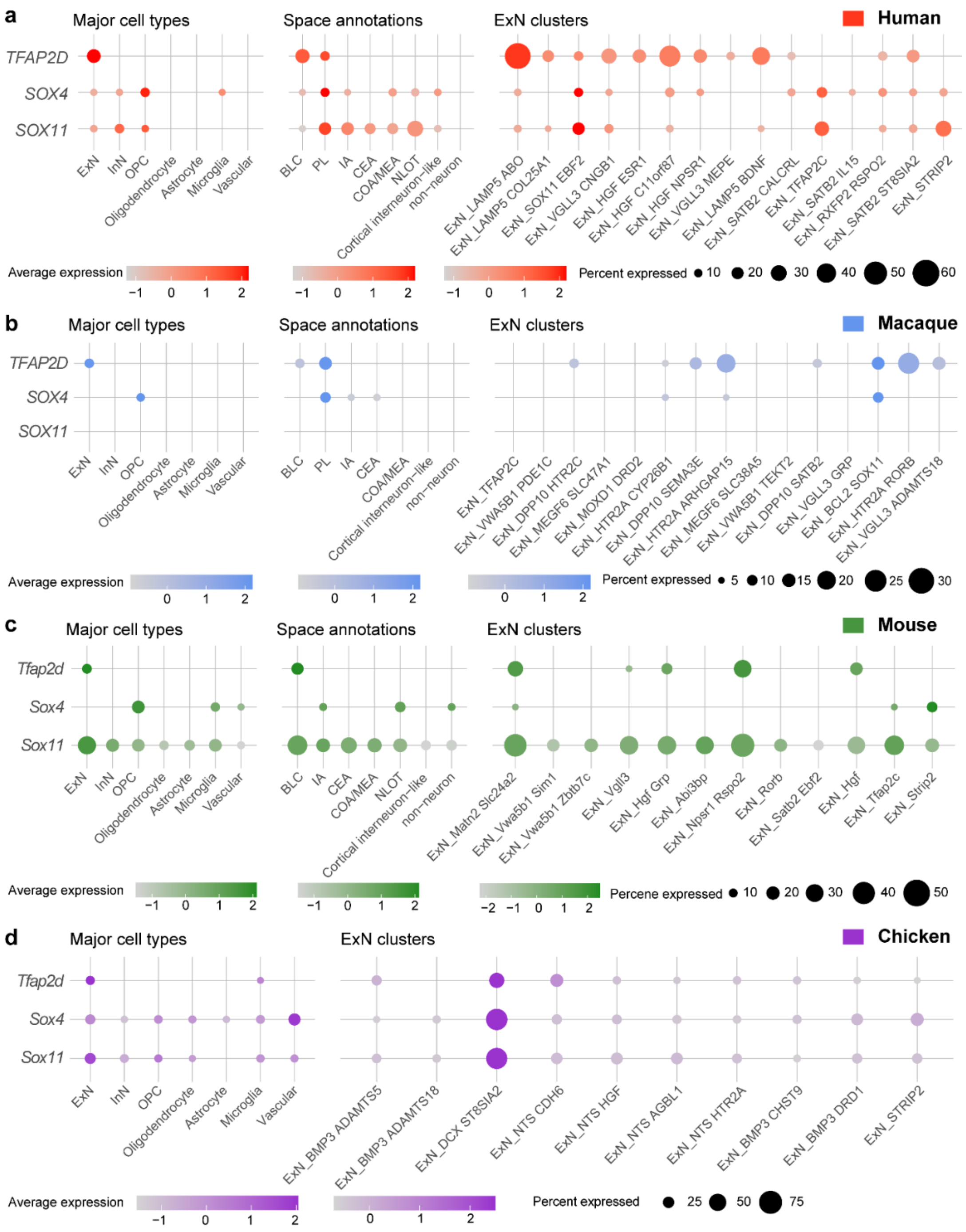


**Extended Data Figure 6. Conserved *Tfap2d* Expression in BLC ExNs Across Mammalian and Non-Mammalian Species**

**a-d**) Dot plot annotations showing the expression of *Sox11*, *Sox4* and *Tfap2d* within 1) the major amygdala cell types in all species, 2) the amygdala nuclei in human, macaque, mouse (space annotations), and 3) ExN clusters within the BLC in human, macaque, mouse and within the caudal dorso-ventral ridge (DVR) of chicken, region homologous to mammalian amygdala [38]. In all species, *Tfap2d* exhibited high expression in ExN within the amygdala. IA, intercalated cell masses; InN, Inhibitory neurons; MEA, medial nucleus of the amygdala; OPC, oligodendrocyte progenitor cells; PL, paralaminar nucleus of the amygdala.

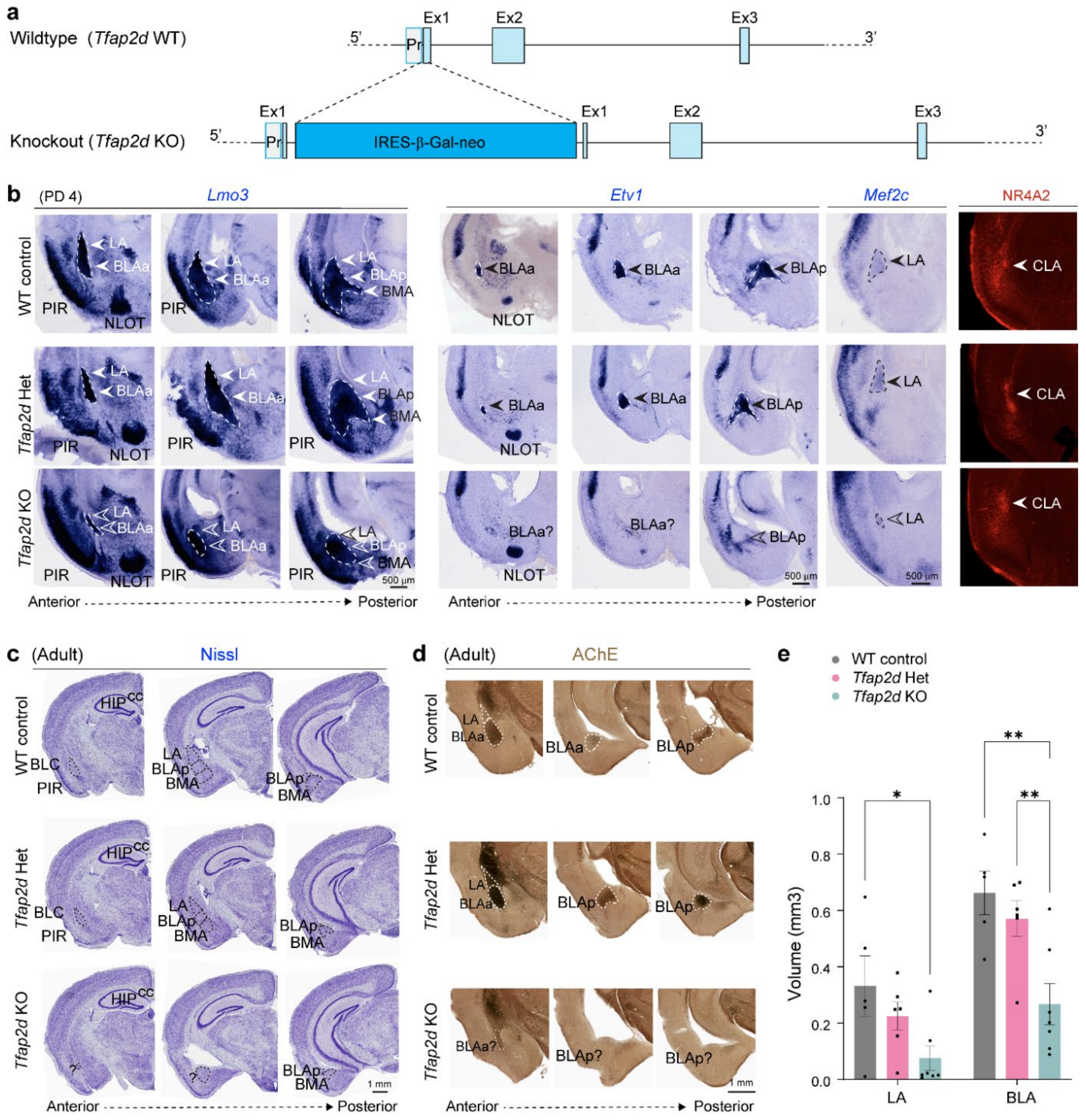


**Extended Data Figure 7. Whole-body Knockout of *Tfap2d* Results in BLC Deficits**

**a**) Schematic depicting the *Tfap2d* wildtype allele and the knockout (KO) allele with the insertion of LacZ cassette within the exon (Ex) 1 of the *Tfap2d* gene locus [30]. **b**) Coronal sections of the ventrolateral cortical regions of the WT control, *Tfap2d* Het and *Tfap2d* KO brains showing the expression of *Lmo3*, *Etv1* and *Mef2c* by *in situ* hybridizations and NR4A2 by immunostaining at PD 4. Reduction in the size of BLC is seen by *Lmo3*, the most affected region is the *Etv1* labelled BLA. **c**) Nissl staining of the coronal sections of the adult WT control, *Tfap2d* Het and *Tfap2d* KO brains. **d**) Coronal sections of the ventrolateral cortical regions of the adult WT control, *Tfap2d* Het and *Tfap2d* KO brains showing the AChE staining used to label the BLA. **e**) Stereological measurements of the volume of LA and BLA in the adult WT control, *Tfap2d* Het and *Tfap2d* KO brains. Mean ± s.e.m. depicted. Two-way ANOVA with Tukey's correction was applied. ** and * represents p values < 0.01, <0.05 respectively. n= 5 (WT), 6 (Het), 7 (KO).

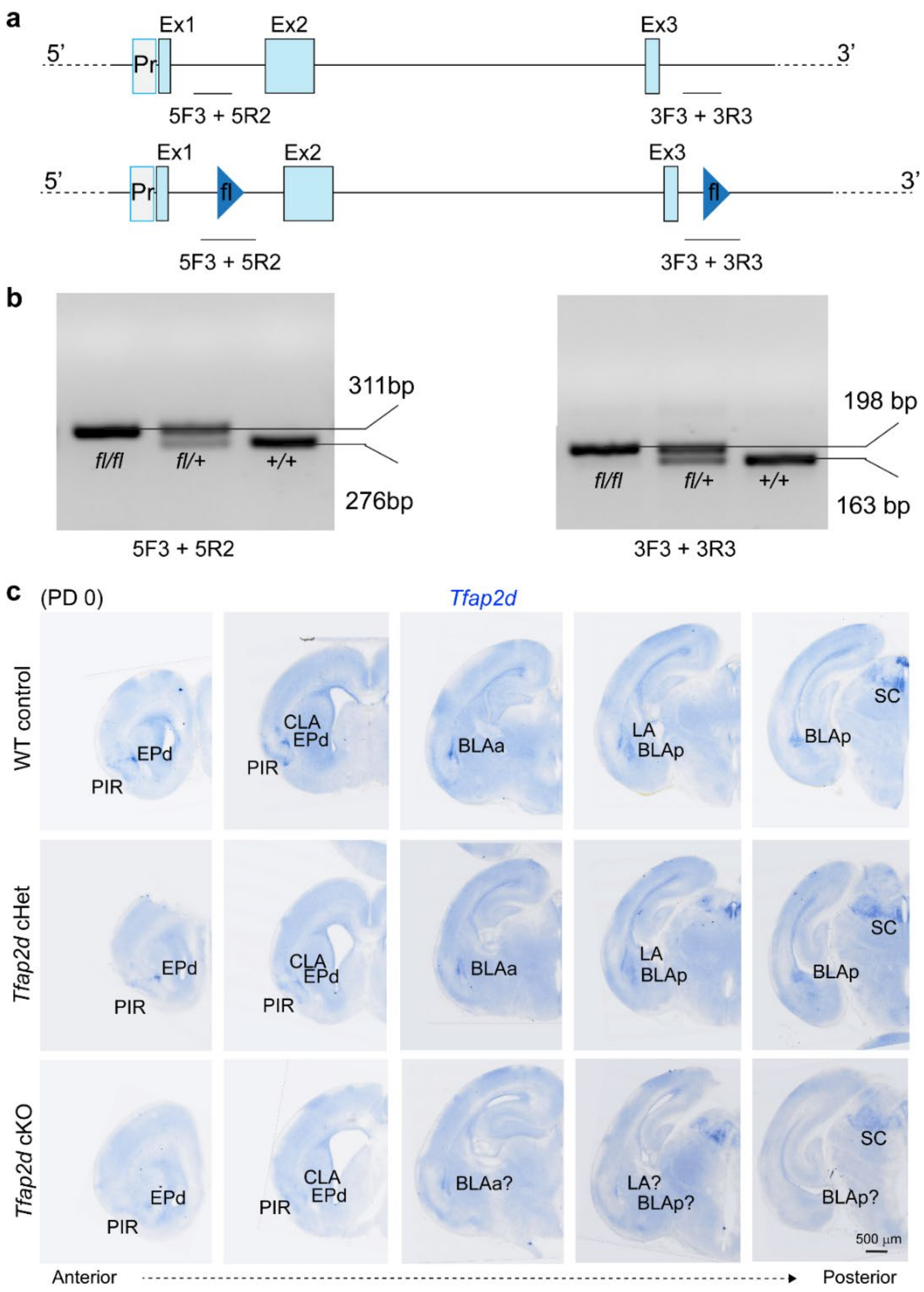


**Extended Data Figure 8. Development of Floxed *Tfap2d* Allele for Conditional Genetic Manipulation**

**a**) Schematic depicting the *Tfap2d* wildtype allele (above) and the floxed allele (below) carrying the insertion of flox (fl) cassettes within intron (In) 1 and In 3 of the *Tfap2d* locus. The regions amplified by PCR using genotyping primers are indicated. **b**) Genotyping agarose gel showing the sizes of the amplified PCR products that can distinguish the fl/fl, fl/+ and +/+ genotypes. **c**) Serial coronal sections of WT control, *Tfap2d* cHet and *Tfap2d* cKO brains at PD 0 showing reduced expression of *Tfap2d* in the ventrolateral pallial structures (CLA, EPd, PIR, BLC) but not in the midbrain (SC, superior colliculus) of cKO mice. Representative images from this panel are also used in Fig. 3a.

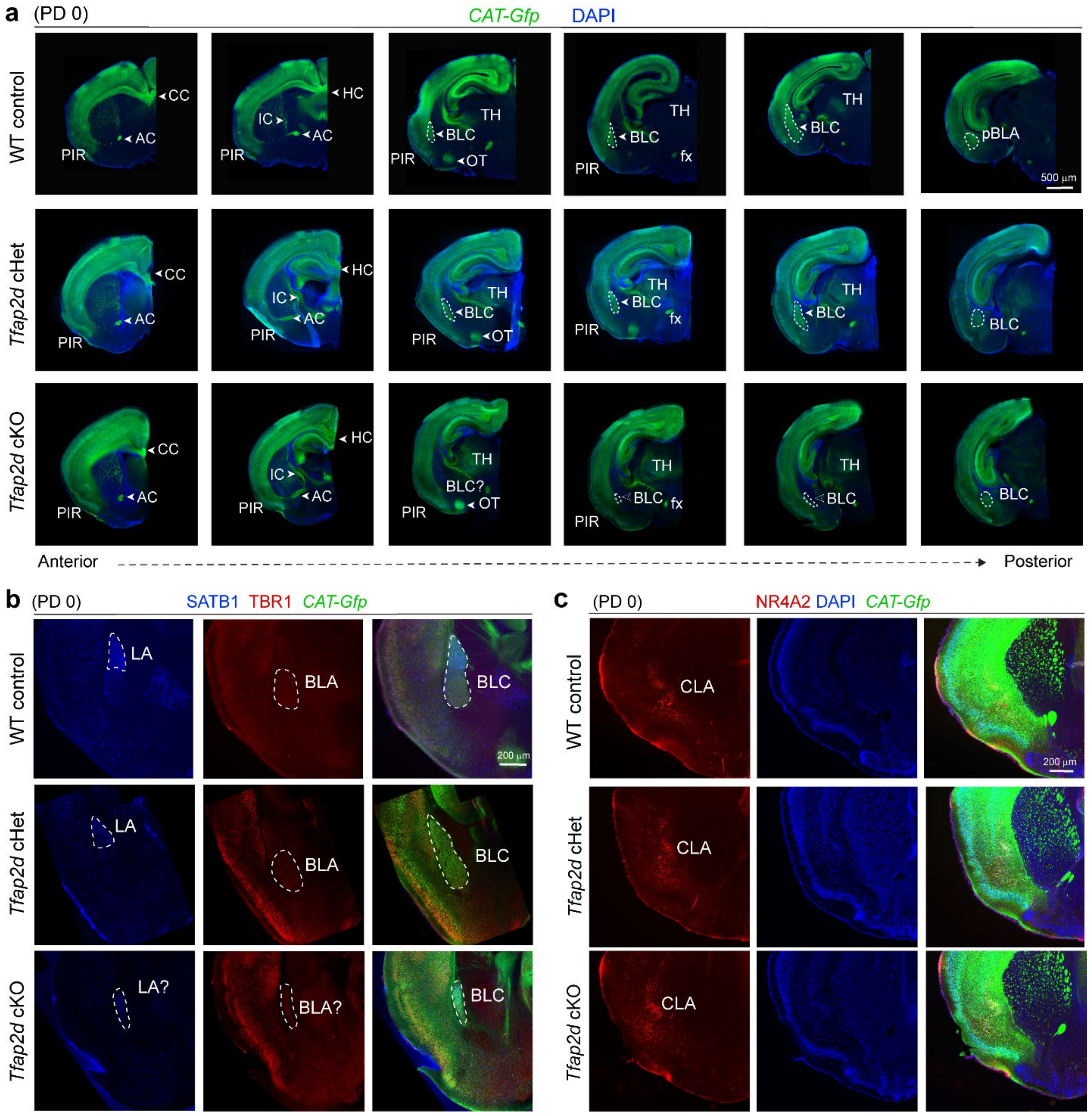


**Extended Data Figure 9. Conditional Post-Mitotic Deletion of *Tfap2d* in Cerebral ExNs Leads to BLC Formation Deficits**

**a**) Serial sections of the WT control, *Tfap2d* cHet and *Tfap2d* cKO brains at PD 0 depicting their gross morphology via visualization of GFP expression driven by the *Neurod6* (*Nex1*)-Cre driver. **b**) Immunostaining showing the expression of TBR1 and SATB1 in the WT control, *Tfap2d* cHet and *Tfap2d* cKO brains at PD 0, highlighting the deficits seen in the BLC. **c**) Immunostaining showing the expression of NR4A2 in the CLA of WT control, *Tfap2d* cHet and *Tfap2d* cKO brains at PD 0. Filled arrows indicate normal morphology whereas open arrows indicated altered BLC structure.

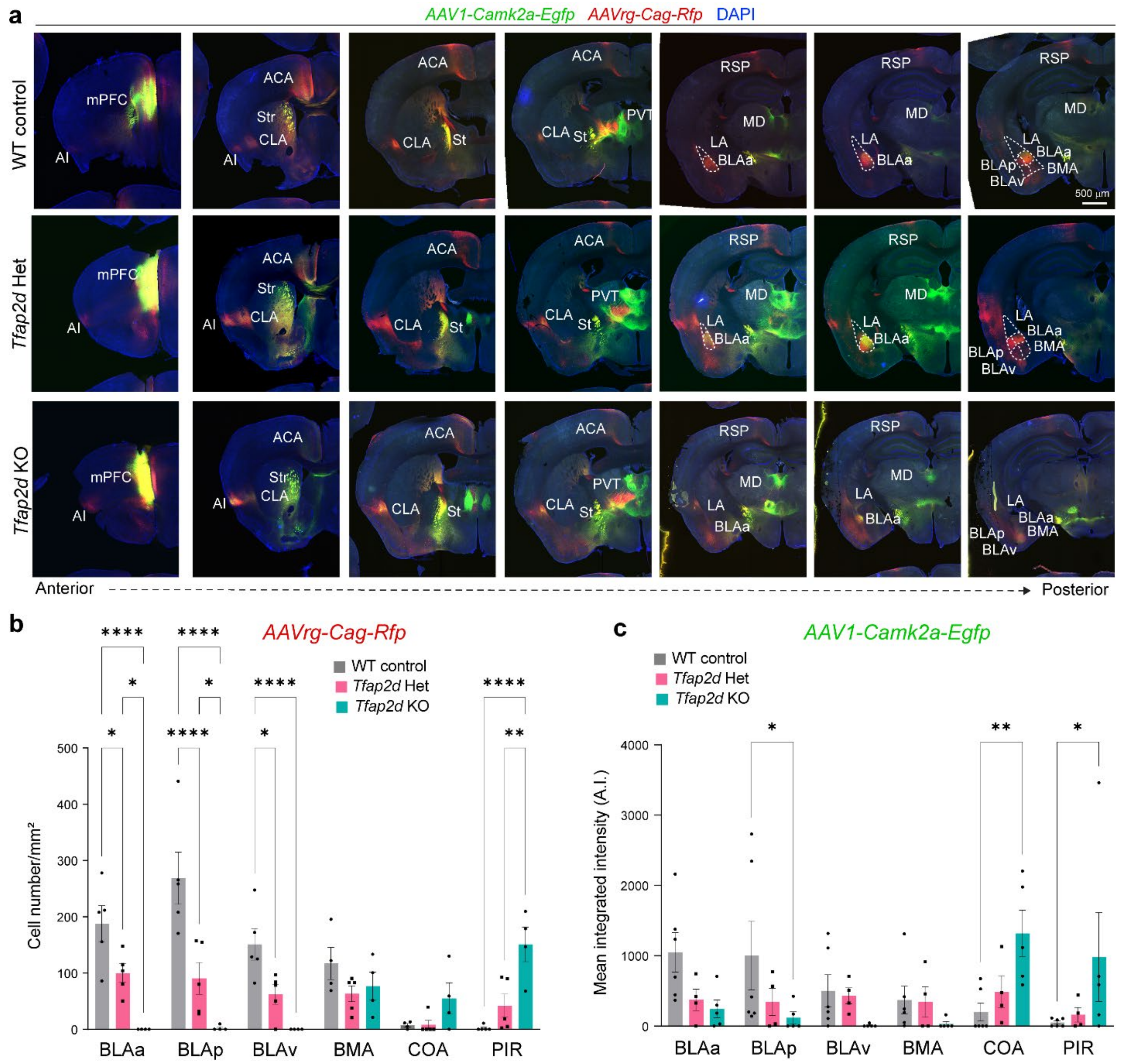


**Extended Data Figure 10.** ***Tfap2d*** **Loss or Haploinsufficiency Impairs BLC-mPFC Connectivity**

**a**) Serial coronal sections of WT control, Tfap2d Het and Tfap2d KO adult brains whose efferent and afferent projections from the mPFC were traced using *AAV1-Camk2a-Egfp* (green; anterograde) and *AAVrg-Cag-Rfp* (red; retrograde), respectively. **b-c**) Bar graphs depicting the number of cells carrying *AAVrg-Cag-Rfp* tracer (b) or mean intensity fluorescence for the axons labelled with *AAV1-Camk2a-Egfp* (c) in various ventrolateral cerebral brain regions of control, *Tfap2d* Het and *Tfap2d* KO adult mice. Mean ± s.e.m. depicted. Two-way ANOVA with Tukey's correction was applied. ****, ** and * represent p values <0.0001, < 0.01, and <0.05, respectively. (n= 5 (WT), 5 (Het), 4 (KO). Str, striatum; St, stria terminalis; PVT, paraventricular thalamus; ACA, anterior cingulate area.

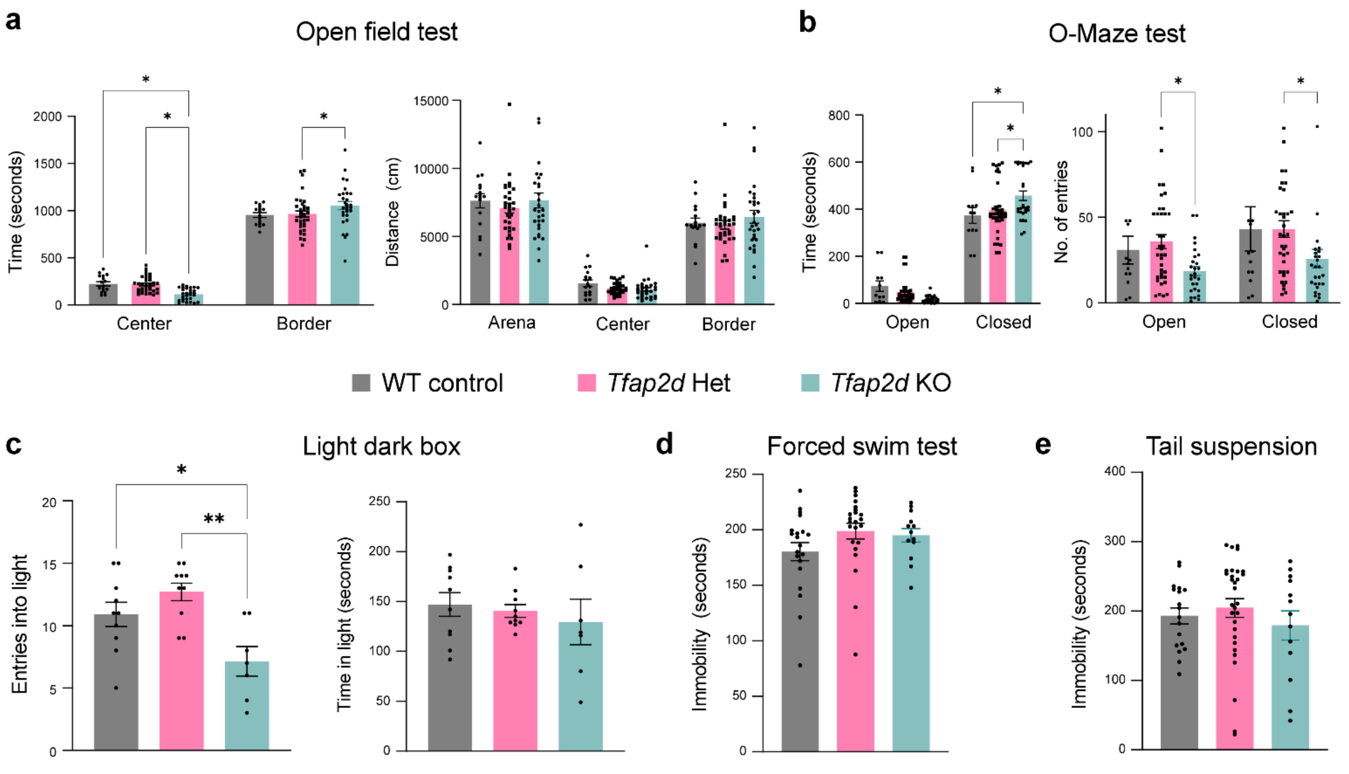


**Extended Data Figure 11. Whole-body Loss or Haploinsufficiency of *Tfap2d* Alters Non-Learned Exploratory Test Behaviors**

**a**) Bar graph showing the cumulative time spent (sec) and distance moved (cm) in the center vs border analysis. *Tfap2d* KO spent significantly more time in the border and moved less in the center, as compared to the WT control, and *Tfap2d* Het. Mean ± s.e.m. depicted. Two-way ANOVA with Tukey's correction was applied. * represents p values, <0.05. n = 16 (WT control), 33 (*Tfap2d* Het), and 29 (*Tfap2d* KO). **b**) Cumulative time spent (seconds) and number (#) of entries in the open arm vs closed arm analysis of the o-maze test. *Tfap2d* KO animals spent significantly more time in the closed arm and had more entries in the closed arm as compared to the WT control and or *Tfap2d* Het animals. Mean ± s.e.m. depicted. Two-way ANOVA with Tukey's correction was applied. ** represents p values <0.05. n = 14 (WT control), 38 (*Tfap2d* Het) ,27 (*Tfap2d* KO). **c**) Number of entries and cumulative amount of time that spent on the light side of the light-dark box. One-way ANOVA with Tukey's correction was applied. ** and * represents p values < 0.01, <0.05 respectively. n = 10 (WT control), 10 (*Tfap2d* Het), and 7 (*Tfap2d* KO). **d**) Cumulative amount of time that animal spent immobile in the forced swim test. One-way ANOVA with Tukey's correction was applied. n = 20 (WT control), 23 (*Tfap2d* Het) and 13 (*Tfap2d* KO). **e**) Cumulative amount of time that animal spent immobile in the tail suspension test. One-way ANOVA with Tukey's correction was applied. n = 18 (WT control), 24 (*Tfap2d* Het), and 13 (*Tfap2d* KO).

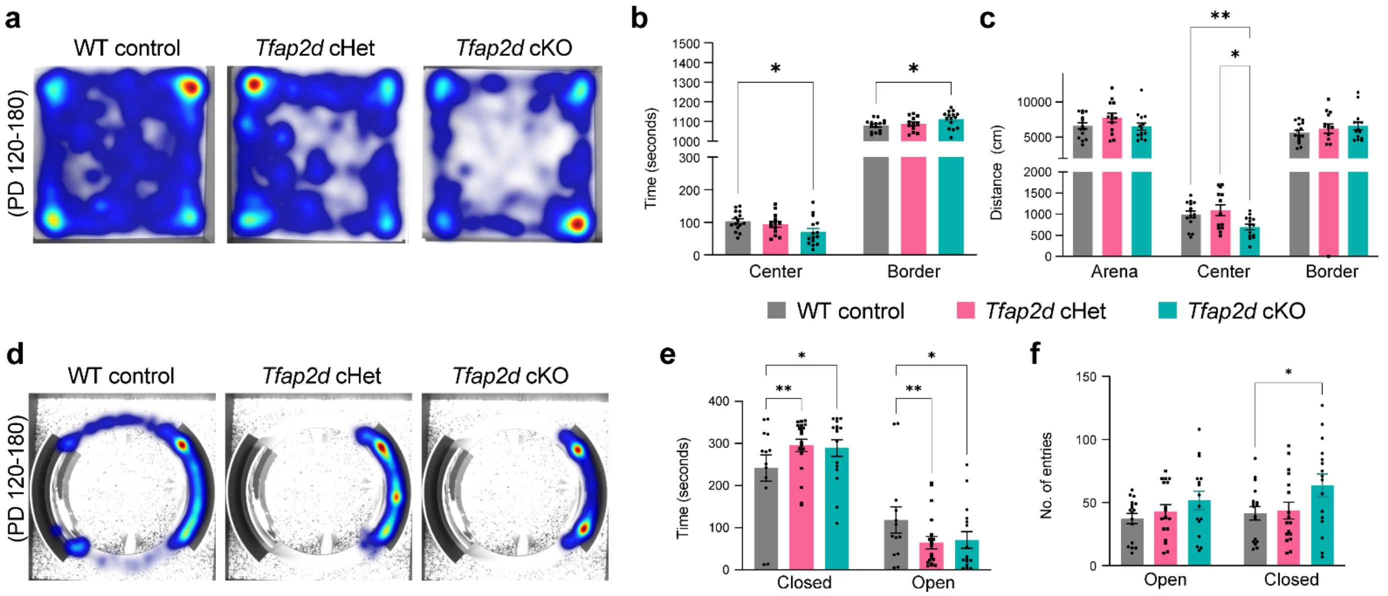


**Extended Data Figure 12. Increased Innate Anxiety-like Behaviors Result from Post-mitotic ExN-specific Deficiency in *Tfap2d***

**a**) Representative heatmaps showing the cumulative movement of WT control, *Tfap2d* cHet and *Tfap2d* cKO animals in the open field test for 20 min. **b-c**) Cumulative time spent (b) and distance moved (c) in the center vs border analysis, showing the *Tfap2d* cKO spent significantly more time in the border and moved less in the center, as compared to the WT control, and *Tfap2d* cHet animals. Mean ± s.e.m. depicted. Two-way ANOVA with Tukey's correction was applied. ** and * represent p values < 0.01, <0.05 respectively. n = 14 (WT control), 14 (*Tfap2d* cHet) and 15 (*Tfap2d* cKO). **d**) Representative heatmap showing the cumulative movement of WT control, *Tfap2d* cHet and *Tfap2d* cKO animals in the o maze test for 6 min. **e-f**) Cumulative time spent and number of entries in the open arm vs closed arm analysis, showing the *Tfap2d* cKO and *Tfap2d* cHet spent significantly more time in the closed arm and *Tfap2d* cKO had more entries in the closed arm as compared to the WT control. Mean ± s.e.m. depicted. Two-way ANOVA with Tukey's correction was applied. ** and * represents p values < 0.01, <0.05 respectively. n = 19 (WT control), 13 (Tfap2*d* cHet) and 16 (*Tfap2d* cKO).

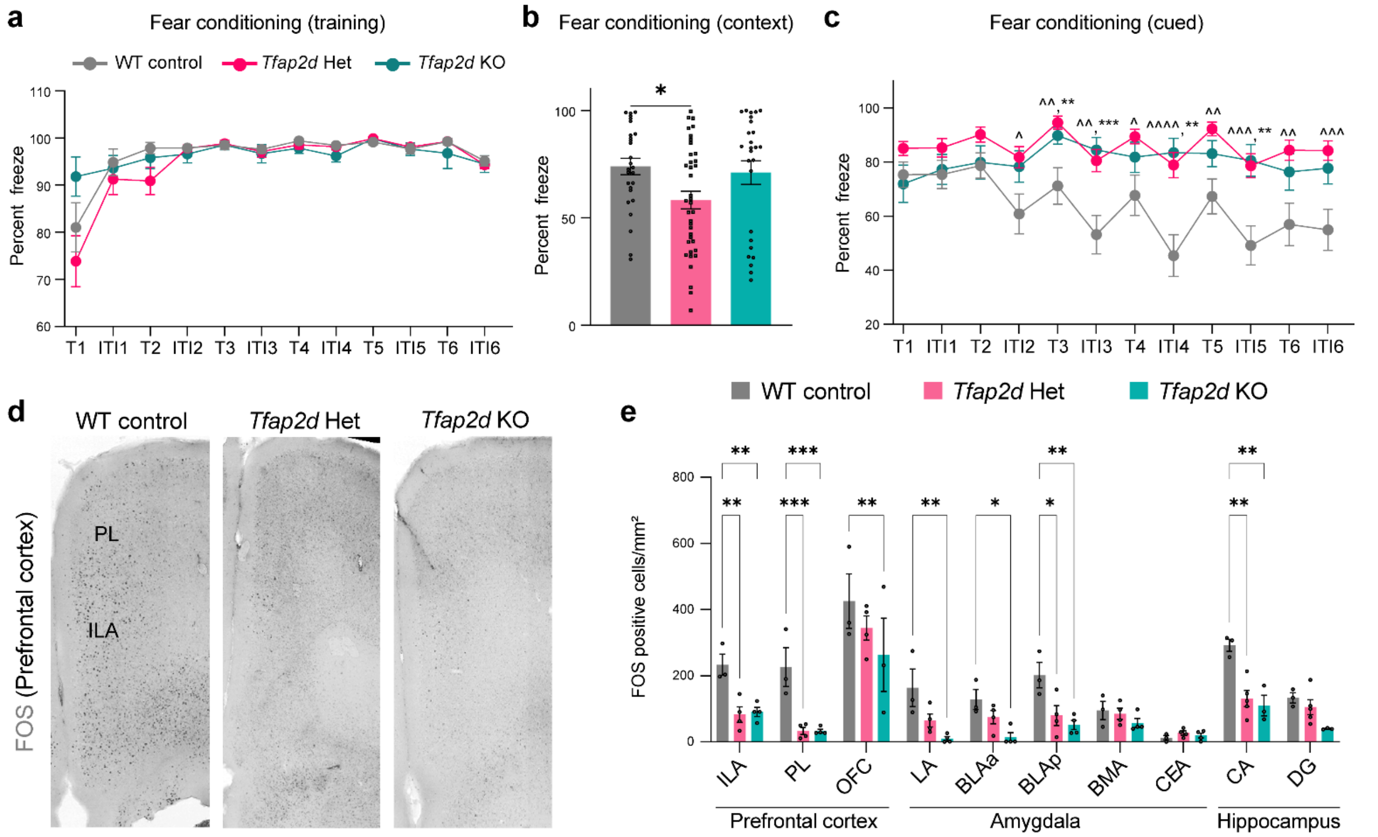


**Extended Data Figure 13. Whole-body *Tfap2d* Loss or Haploinsufficiency Alters Threat Response in the Fear Conditioning Test**

**a**) Line plot showing that during learning phase WT control, *Tfap2d* Het and *Tfap2d* KO mice freeze for same amount of time by tone 6. Mean ± s.e.m. depicted. Two-way ANOVA with Tukey's correction was applied. **b**) Bar plot showing conditioned responses of the WT control, *Tfap2d* cHet and *Tfap2d* cKO mice as percent of time the animals froze while in the training context. Mean ± s.e.m. depicted. One-way ANOVA with Tukey's correction for multiple comparisons was applied. * represent p values <0.05. **c**) Line plot showing cued responses of the WT control, *Tfap2d* cHet and *Tfap2d* cKO mice. Mean ± s.e.m. depicted. Two-way ANOVA with Tukey's correction was applied. *** and ** represent p values < 0.001, <0.01 respectively for WT vs cKO and ^^^^, ^^^ and ^^ represent p values < 0.0001, <0.001 and <0.01 respectively for WT vs cHET. n = 27 (WT control), 38 (*Tfap2d* Het), 26 (*Tfap2d* KO). **d**) Representative image of the mPFC showing immunostaining for FOS performed on the WT control, *Tfap2d* cHet and *Tfap2d* cKO brains isolated after 60-90 min of cued memory test. **e**) Graph depicting the number of FOS positive cells per mm$^2$ detected within the different brain regions of the WT control, *Tfap2d* cHet and *Tfap2d* cKO brains isolated after 60-90 min of cued test. Mean ± s.e.m. depicted. Two-way ANOVA with Tukey's correction was applied. ***, **, * represents p values < 0.001, <0.01, <0.05 respectively.

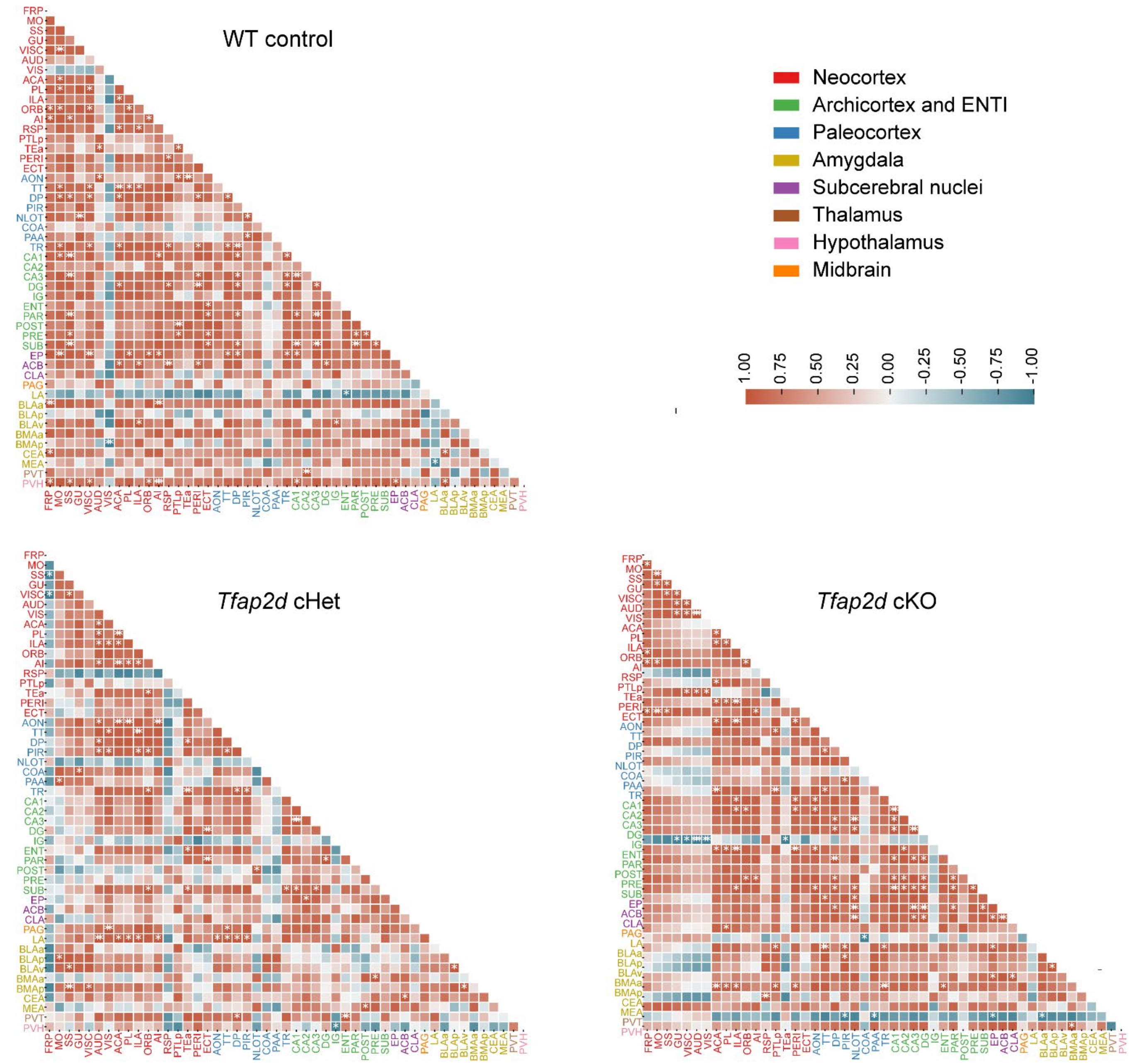


**Extended Data Figure 14. Alterations in Functional Connectivity Following Cued Testing Due to Post-mitotic ExN-specific Deficiency in *Tfap2d***

Pearson correlation heatmap showing the functional connectivity between different brain regions measured by the FOS immunostaining of the WT control, *Tfap2d* cHet and *Tfap2d* cKO mice after the cued test. ***, **, * represents p values < 0.001, <0.01, <0.05, respectively (n= 4/ genotype).

**List of Tables:**

**Extended Data Table 1: Genes with Increased Expression Exclusively in the *Sox4; Sox11* cdKO as Compared to *Sox4* cKO, *Sox11* cKO and Controls.**

**Extended Data Table 2: Genes with Decreased Expression Exclusively in the *Sox4; Sox11* cdKO as Compared to *Sox4* cKO, *Sox11* cKO and Controls.**

**Extended Data Table 3: List of the Oligos and Primers.**

**Extended Data Table 4: List of Abbreviations for Figure 5d-e and Extended Data Figure 14a.**

**Extended Data Table 5: List of Regions and Abbreviations for Extended Data Figure 18.**

### Methods

#### Animals

All experiments were carried out with the protocol approved by the Committee on Animal Research at Yale University. The research methodologies for the utilization of mice (Mus musculus) and rhesus macaques (Macaca mulatta) were implemented following the guidelines sanctioned by the Yale University Institutional Animal Care and Use Committee (IACUC) and the directives of the US National Institutes of Health. The care and management of the animals were conducted within the precincts of the Yale Animal Resource Center, ensuring controlled environments for both prenatal and postnatal development stages of mouse and primate specimens. The mice were group-housed, maintaining a density of less than five individuals per cage, under environmental conditions regulated at 25°C and 56% relative humidity, complemented by a photoperiod consisting of 12 hours of light and 12 hours of darkness. Nutritional needs were met *ad libitum*, coupled with veterinary oversight provided by the center's staff. The genetic lineage of the subjects was maintained on a C57BL/6J strain, with experimental cohorts comprising both genders, randomly designated to respective studies. The detection of a vaginal plug in the murine subjects was recorded as gestational day 0.5, marking the initiation of the experimental timeline. The *Tfap2d*-Lacz mice were a kind gift from Markus Moser and the generation and genotyping were previously described (Extended Data Fig. 7a; [30]). These mice were generated by inserting a Lacz cassette that disrupts the Exon 1 of the Tfap2d locus, abolishing the proper expression of the trapped allele, Tfap2d. The *Nex1-Cre* (*Neurod6-Cre*) were a kind gift from Klaus-Armin Nave lab and *CAT-Gfp* mice were procured from Jackson's lab [51]. Please refer to Extended Data Table 3 for genotyping details.

#### Postmortem Human and Macaque Brain Tissue

Human brain samples were collected postmortem at 17 post conception weeks. Rhesus macaque brain samples were collected postmortem at 105 postconceptional days. Whole slabs or whole hemispheres were postfixed in 4% PFA for 48 hours and then cryoprotected in an ascending gradient of sucrose (10%, 20%, 30%). Tissue was handled in accordance with ethical guidelines and regulations for the research use of human brain tissue set forth by the NIH (http://bioethics.od.nih.gov/humantissue.html) and the WMA Declaration of Helsinki (http://www.wma.net/en/30publications/10policies/b3/index.html). All experiments using non-human primates were carried out in accordance with a protocol approved by Yale University's Committee on Animal Research and NIH guidelines.

#### Generation of *Tfap2d* Floxed Mice

Guide RNA sequences (gRNAs) to insert *flox* sites were designed using an online program (http://zlab.bio/guide-design-resources) [52]. gRNAs with the minimum off-target effects were selected, and DNA oligos carrying guide RNA sequences were cloned into pX330 vector. The sequences of the guide oligos used are listed in Extended Data Table 3. Cas9 mRNA were in vitro transcribed from vector px330 linearized by restriction enzyme NotI-HF (New England Biolabs, #R3189S) and purified by phenol/chloroform extraction. We inserted the flox sites into intron 1 and intron 3. Founders were genotyped and bred at least 3 generations to exclude chimeras. Genotyping of mice for floxed allele was performed using primers listed in Extended Data Table 3, Extended Data Fig. S8 a-b.

#### *In situ* hybridization

The cDNA (*Tfap2d* human: ENSG00000008197; *Tfap2d* mouse:; *Tfap2d* macaque: ENSMMUT00000004057; Lmo3: ENSMUSG00000030226; Etv1: ENSMUST00000095767; Mef2c: ENSMUST00000163888, Cyp26b1: ENSMUST00000077705) for generating the human and mice probes was procured from Dharmacon and the plasmid was cut using NOT1HF enzyme. For the *Tfap2d* mice probe three different probes were amplied from PD 0 and adult cDNA using the primers listed in Table S3. The *Tfap2d* V3 probe worked best across ages and was used in the manuscript for detection of *Tfap2d* expression. The macaque probe was generated using respective neocortical tissues cDNA as a template by TA-cloning kit (Invitrogen, K202020). The cDNA clones were purified through the Qiagen clean up kit (Qiagen, 28104) and *in vitro* transcription (Millipore Sigma- Roche, 10999644001) was performed per the manufacturer's instructions. Templates were purified by phenol/chloroform extraction and Digoxigenin labelled probes were synthetized using T3 (Roche, RPOLT3-RO) and T7 RNA polymerases (Roche, RPOLT7-RO) respectively, and RNA labelling mix (Roche, 11277073910) according to the manufacturer's instructions. Probes were purified by ethanol precipitation, quantified, quality controlled, and stored at -80°C until hybridization. For *in situ* hybridization (ISH), slide-mounted cryo-sections at 60-70µm thickness were processed. Briefly, brains were fixed overnight at 4°C in 4% PFA (Electron Microscopy Sciences, 15711) diluted in Dulbecco's phosphate buffer (DPBS) (Thermo FisherScientific, 14190144), equilibrated for 12 hours at 4°C in 10% sucrose, and another 12 hours at 4°C in 30% Sucrose in DPBS. Fixed brains were then embedded in OCT (Scigen, 23-730-571) and sliced on a cryostat (Leica Biosystem, CM1800). Slides were stored at -80°C until processed for in situ hybridization. Sections were first post-fixed in 4% PFA in PBS for 15 minutes at room temperature, washed with PBS, and treated with 0.5 µg/ml proteinase K solution (15 min at RT for P0 and 30 at RT min for adults). Slides were post fixed in 4% PFA in PBS for 15 minutes at room temperature, washed with PBS. Slides were treated with acetic acid and triethonalamine solution for 10 mins, followed by PBS washes. Slides were then submerged in hybridization buffer (5X SSC, 50% Formamide, 20% SDS and 250 µg/ml of Torula yeast RNA, aBSA (25mg/ml), Heparin Stock (50mg/ml)) supplemented with 1000ng/ml of the appropriate digoxigenin-labeled probe at 70°C overnight. Sections were washed three times for 45 minutes at 70°C in 2X SSC, 50% formamide, 1%SDS, and then three times with TBST and incubated overnight at 4°C with an anti-digoxigenin antibody conjugated to alkaline phosphatase (1:5000, Roche, 11093274910). Sections were washed and then rinsed in the substrate buffer (100mM Tris-Cl pH9.5, 100mM NaCl, 50mM MgCl2, 0.1% Tween-20) before being overlaid with NBT/BCIP substrate (Roche). Color development was done at room temperature in the dark until the desired signal was reached. Finally, sections were rinsed in EDTA solution and DPBS, washed in water and mounted with VectaMount® AQ Aqueous Mounting Medium (Vector Laboratories, H-5501-60).

#### RNA In Situ Hybridization (RNAscope) Analysis on Chicken Brain Tissue

The brains from the embryonic chicken were dropped fixed in 4% PFA overnight at 4°C. They were sectioned at 20 µm and slides were stored in -80°C until use. RNAscope was performed as per the multiplex fluorescence protocol from ACD (Advanced Cell Diagnostics) bio with slight modification. Briefly, the slides were thawed and washed with 1X PBS for 5 mins at RT, followed by incubation at 60°C for 30 min in the HybEZ hybridization system (321711). Slides were fixed for 30 min at 4°C, followed by dehydration with ethanol (50%, 70%, and 100%, each for 5 mins). Slides were then airdried for 5 mins and then incubated at 60°C for 15 min. Slides were treated with hydrogen peroxide for 10 min and washed with distilled water twice for 2 mins. They were again incubated at 60°C for 15 min and then washed with boiling water for 10sec. Further, target retrieval is performed using the RNAscope Target Retrieval reagents for 5 min in a boiling beaker. The samples were washed with distilled water again and then incubated with 100% ethanol for 5 min. Slides were incubated at 60°C for 15 min. Using a barrier pen a boundary is created and the slides are incubated with 5 drops of the RNAscope protease III for 30 mins. Samples are then washed with distilled water and incubated with the *Tfap2d* probe for 2 h at 40°C. After that samples were incubated sequentially with RNAscope Multiplex FL v2 Amp 1, RNAscope Multiplex FL v2 Amp 2, and RNAscope Multiplex FL v2 Amp 3 for 30 min each with intermittent washes with 1X wash buffer twice for 2 min. The slides were then incubated with HRP-C3, followed by TSA amplification, dehydration and mounting. Slides were imaged on the VS 200 microscope (Olympus Microscopy).

**Immunohistochemistry**

The brains isolated from prenatal and early postnatal animals were fixed overnight in paraformaldehyde (PFA) at 4°C. Immunostaining was performed on the sections of 60-70 um thickness cut on vibratome. Sections were blocked for 1hr at RT with blocking buffer consisting of 10% Donkey serum (Jackson ImmunoResearch, AB_2337258), 1%BSA (Millipore Sigma, A4612), 0.3% Triton X-100 (which one) in 1X PBS. After blocking, sections were incubated overnight at 4°C with primary antibodies for SATB1 (Santacruz, sc-5989) , TBR1 (Abcam, ab183032) , NR4A2 (R&D systems, AF2156), CUX1 (Santacruz, sc-13024), BCL11B (Abcam, ab18465), FOS( Cell Signalling, 2250) GFP (1:500; Abcam, ab13970), and RFP (1:500, Abcam, ab124754). Sections were washed with washing buffer (0.3% Triton X-100 in 1X PBS) thrice, 10 min each and were incubated with secondary antibodies (Jackson ImmunoResearch ) for 1 hr at RT, followed by three washes with washing buffer for 10 min each. Sections were mounted onto glass slides with vector shield (Vector labs, H-1000) and sealed with nail-polish. All the slides were stored at -20°C for further analyses. Antigen retrieval was performed on the brain sections prior to NR4A2 and TBR1 immunostaining (Dako Antigen Retrieval Solution, GV80511-2). For acquiring the images, LSM 800 (Zeiss Microsystems) and VS 200 microscope (Olympus Microscopy) were used. Analyses of the images was done using ZEN software, Qupath (Bankhead et al 29203879), OlyVIS or Fiji (Schindelin et al 22743772) using BioFormats plugin.

**Generation and Analysis of RNA-seq Data**

The cerebral cortex and amygdala from P0 Sox4; Sox11 cdKO, Sox11 cKO, Sox4 cKO and littermate control pups were collected (n = 3). Total mRNA was extracted using mirVANA Kit and DNAse digestion was performed using Tubro DNAse. Libraries were prepared with Illumina TruSeq mRNA preparation kit (Illumina RS-122-2101) for whole cortices, as per the manufacturer's instructions. Libraries were quality controlled by Tapestation/ Bioanalyzer analysis and sequenced on the Illumina HiSeq 2000 platform at Yale Center for Genome Analysis (YCGA) to generate 75 bp single reads. Sequencing data were quality controlled by FastQC and aligned to the mouse genome (NCBI37/mm9) using STAR (v2.4.0e) (https://doi.org/10.1093/bioinformatics/bts635). To improve the mapping quality of splice junction reads, mouse gene annotation retrieved from the GENCODE project (version M1) was additionally provided (https://doi.org/10. 1101/gr.135350.111). Command line ‘‘–sjdbOverhang 74’’ was used to construct a splice junction library. At least 10 million uniquely mapped reads were obtained for each sample. Differential gene expression DEX analysis was performed by the R package DESeq2 (https://doi.org/10.1186/gb-2010-11-10-r106) and PCA analysis was performed by the R package prcomp. Genes with $|\log_2\text{FoldChange}| \geq 0.5$ and a false discovery rate (FDR) < 0.01 were classified as DEX. Integrated analysis was performed to identify the number of common or distinct DEX between *Sox4*; *Sox11* cdKO, *Sox11* cKO and *Sox4* cKO.

928 up- and 659 down- uniquely expressed genes in *Sox4*; *Sox11* cdKO were selected as candidate downstream targets, as shown in Extended Data Table S1 and S2.

We further searched for cell types of amygdala using the public scRNA-seq dataset [36]. To find cell type marker genes, we used the Seurat *FindMarkers* function. We performed the Wilcoxon Rank Sum test, and considered genes with a minimum expression ratio of 0.2, adjusted p-value less than 0.05, *logfc.threshold* greater than 0.25, expression percentages less than 0.2 in other subtypes, and a foldchange of expression percentages greater than 1.5 between the Excit_B subtype and the second subtype where the genes are detected as marker genes.

**Human Gene Expression Analysis Utilizing the Allen Brain Atlas and the BrainSpan Atlas**

The images of brain structures presented in Figure 2d were generated utilizing the Freesurfer software [53]. To identify cortical regions, the Desikan-Killiany atlas was employed, a commonly used atlas for human brain mapping [54]. TFAP2D expression values from two probes (A_23_P386973, CUST_2289_PI416261804) were taken from the Allen Brain Atlas[34] (adult data) and summed. Then, the average of $\log_2$ expression values across six distinct individuals was computed. To visualize the expression levels of the gene, plot data, and to establish correspondence between the Desikan-Killiany atlas and the Allen Brain atlas probes, the methods described in a previous study were used [55]. Subcortical regions were visualized using data from the Allen Brain Atlas website (http://atlas.brain-map.org). A standardized $\log_2$ gene expression color bar graph across both cortical and subcortical regions was created using a custom R script developed in-house. To generate the graphs in Fig 2c (Allen Brain Atlas adult data[34]) and Extended Data Fig. 4a (BrainSpan Atlas data[35], midfetal data; www.brainspan.org), the $\log_2$ TFAP2D expression values of the two different probes were averaged within a sample. For Fig 2c, top level structure names were changed to follow cortical cytoarchitectonics designations.

**Comparative Analysis of *Tfap2d* Gene Expression Across Mammalian and Non-Mammalian Species**

To assess the expression patterns of *TFAP2D* across different species, we reanalyzed public single-nucleus transcriptome datasets for amygdala in human (Homo sapiens), macaque (Macaca mulatta), mouse (Mus musculus), and chicken (Gallus gallus). We checked the expression level of *Tfap2d*, *Sox4*, *Sox11*, *Lmo3*, *Etv1* and *Mef2c* across major cell types, space clusters, and cell type clusters, as identified by the original paper [38]. For cross-species comparison, we only included the excitatory neurons.

**Analysis of *Tfap2d* Regulatory Regions and SOX11 ChIP-seq Data**

To determine what regions within the *Tfap2d* locus can be potential enhancers, we re-processed a single-cell ATAC-seq embryonic dataset of the mouse cortex [33]using Signac version 1.1.0 (https://satijalab.org/signac/), to obtain ATAC-seq peaks that are accessible in migrating ExNs in PCD 13.5 onwards. We further refined the list of putative peaks by requiring that they intersect with occurrences of *Sox4* or *Sox11* JASPAR motifs (MA0867.1, MA0867.2, MA0869.1, MA0869.2). Similarly to regions directly bound by SOX4 and SOX11 within *Tfap2d* locus we analyzed data generated from ChIP-seq using an antibody against H3K27ac, a histone mark enriched in promoters and enhancers [32]. The SOX11 ChIP-seq data was directly accessed from the Bergsland et al 2011[24].

**Chromatin Immunoprecipitation and RT-PCR**

Cortices from PCD 15.5 mice were isolated and were crosslinked with a formaldehyde solution at a final concentration of 1% at room temperature for 10 minutes. L-Glycine (AmericanBio) at a final concentration of 125mM was added and incubated for 5 minutes at room temperature to quench the cross-linking. The tissue was spun down and lysed in the hypotonic solution (50 mM Tris-Cl pH 7.5, 0.5% NP40, 0.25% Sodium Deoxycholate, 0.1% SDS, 150 mM NaCl) on ice for 10 minutes to obtain the nuclei. Nuclei were centrifuged at 600g for 5 minutes at 4°C and pellets were resuspended in the SDS lysis buffer (1% SDS, 10mM EDTA and 50mM Tris-Cl, pH 8.1) prior to being sheared into 200-500 bp size fragments using a sonicator (M220 Focused-ultrasonicator, Covaris). The sheared DNA was diluted with the ChIP dilution buffer (0.01% SDS, 1.1% Triton X- 100, 1.2mM EDTA, 16.7mM Tris-Cl, pH 8.1, 167mM NaCl) and was precleared with magnetic protein A/G beads (Thermo Scientific) for 1 hour at 4ºC. For ChIP, 10µg anti-SOX11 (Abcam, ab229185), 1 µg PolII (Sigma-Aldrich, 05-623) and 10µg of IgG (Diagenode, C15410206) was used. Samples were incubated on constant rotation overnight at 4°C. Magnetic protein A/G beads (EMD Millipore,166-63) were blocked with 1mg/ml bovine serum albumin (BSA) (Sigma-Aldrich) and tRNAs were added to the chromatin-antibody complexes for 4 hours at 4°C. Beads were washed with low salt (0.1% SDS, 1% Triton X-100, 2 mM EDTA, 20 mM Tris-Cl, pH 8.1, 150 mM NaCl), high salt (0.1% SDS, 1% Triton X-100, 2 mM EDTA, 20 mM Tris-Cl, pH 8.1, 500 mM NaCl), LiCl (0.25 M LiCl, 1% IGEPAL CA630, 1% deoxycholic acid (sodium salt), 1 mM EDTA, 10 mM Tris-Cl, pH 8.1), and TE (AmericanBio), sequentially for 3 minutes each. ChIP DNA was incubated overnight at 65°C for reverse cross-linking and subjected to RNAse A treatment (37°C, 1hour) and Proteinase K treatment (55°C, 2 hours), then purified on PCR purification columns. For input control, 5 µg cross-linked chromatin from each sample was also treated by reverse cross-linking, RNAse A (Thermo Scientific), and Proteinase K (Millipore Sigma) together with IP samples and purified by PCR purification columns. DNA amounts were quantified by PicoGreen assay (Thermo Scientific). The samples were eluted in 20 µl of TE. Each sample is diluted 1:5 and processed for RT-PCR using the BioRad Sybr green mix and BioRad machine. Samples were run in triplicates and fold enrichment was calculated over the input. Two-way repeated measures ANOVA with Tukey's multiple comparison correction was applied.

#### Nissl Staining

Perfused brains were sectioned at 40 µm thickness and mounted and dried on slides. A 1:12 series through the brain was stained for Nissl. Briefly, sections went through a demyelination step including first an ascending series of ethanol followed by a descending series of ethanol, stained with Cresyl violet acetate stain for 5 minutes and then briefly washed with water and ethanol. Sections were destained using a mild differentiating solution (50% ethanol + 5 drops of glacial acetic acid) and further dehydrated in an ascending series of ethanol before being placed in xylene and coverslipped. Slides were scanned with the Leica Aperio digital slide scanner at 20x magnification. Samples were observed for gross morphological alterations due to genotype.

#### Acetylcholinerase Histochemistry and Stereological Analysis

For each animal, a 1:12 series of sections through the brain was stained with acetylcholinerase [56]which demarcates the La from the BLA. Each series contained between four and six sections through the anterior-posterior extent of the amygdala. Slides were scanned with the Leica Aperio digital slide scanner (Leica, Wetzlar, Germany) at 20x magnification. Images used for stereological estimates were cropped using Leica's Aperio ImageScope software and saved as JPEG. The Cavalieri estimator probe, implemented in

Stereoinvestigator (MBF Bioscience, Williston, VT USA) was used to calculate BLA and La volumes using a grid size of 10 μm. Volume estimates used are corrected for overprojection.

**Diffusion Tensor Imaging (DTI)**

Nine postmortem mouse brains (3 WT control, 3 *Tfap2d* Het, 3 *Tfap2d* KO) were perfusion-fixed in with 4% paraformaldehyde solution in 0.1M PBS for 24 hr. Following perfusion-fixation, mouse brains were immersed and stored in 0.1M PBS for 24 hours and were transferred to Fomblin (SPI Supplies 69991-67-9) just before imaging. Diffusion MRI was acquired on a BioSpin 9.4T MRI (Bruker) machine for each subject using a 3D echo-planner-imaging (EPI) diffusion sequence (b=1500s/mm2; 30 directions) with the following parameters[57]: repetition time=1250ms; echo time=26ms. The resolution was 0.1mm isotropic. Overall scanning time was 19.5h.

**DTI Image Processing and Tractography**

Cerebral cortical and subcortical regions of interest (ROI) and thalamus were manually defined according to Paxinos [58] and the Allen Mouse Brain Atlas [59] (https://mouse.brain-map.org/static/atlas) by D.A., N.S., and N.K. without prior knowledge of the experimental groups. Whole-brain deterministic tractography[60-62] was performed with Diffusion Toolkit (Version 0.6.4.1) software with a fractional anisotropy threshold of 0.02 and an angular threshold of 60 degrees. Fibers that pass through each pair of ROIs were delineated and quantified. Visualization of the tracts and mouse brain was generated using MRtrix361 software (v.3.0.0-65-g91788533). Circular visualization of the tractography derived connectivity between brain regions was generated using Circos software package [63].

**Retrograde and Anterograde Neuronal Tracing**

To determine the effects of BLC deficits associated with *Tfap2d* loss and confirm the DTI results, we performed the AAV based tracing from the mPFC, with retrograde and anterograde tracers injected simultaneously into the mPFC of PD 120-180 animals. In brief, animals were anaesthetized by injecting a ketamine/xylazine solution and head fixed in a stereotactic frame. Thirty minutes before surgery, buprenorphine was administered. After lubricating the eyes and shaving the fur, an incision of <1 mm was made. A craniotomy was made with a round 0.5-mm drill bit at the desired coordinates (for the mPFC: medial–lateral±0.35, anterior–posterior 2.0, dorsal–ventral 2.5). Using a Hamilton neuros syringe (0.5 ml), we mixed and injected 100 nl of each of *AAVrg-Cag-Tdt* (59462-AAVrg, Addgene) of *AAV1-Camk2a -eGfp* (50469-AAV1, Addgene) into the mPFC. To prevent the virus from spreading along the injection tract, the needle was held in place for at least 10 min. After injections, the skin was sutured, and the animals were returned to the cage. Approximately 3 weeks later, the animals were euthanized, and their brains were collected. The brains were coronally sectioned on a vibratome to obtain 70-μm-thick sections. After staining the sections with anti-GFP antibody (1:500; Abcam, ab13970) and anti-RFP antibodies (1:500, Abcam, ab124754), the sections were imaged with VS 200 (Olympus Microscopy). The images were analyzed using the Qupath software where the outline for the ROI was created and the number of cells projecting to mPFC were counted above threshold and the intensity of the axons in each region was measured using the Image J plugin.

**Mouse Behavioral Assessments**

We performed a battery of tests on the mice to determine their exploratory, innate, or learned anxiety-related behaviors[64]. The experimental cohort comprised of male and female littermates' aged between PD 120-180. Animals were acclimatized in the room for 30 mins to 1hr before the beginning of any behavior experiments. The experiment began with the apparatus being sanitized using 70% ethanol. In the following trials, water was used for cleaning instead. The first trial featured control animals and to mitigate the strong ethanol scent that might alter their behavior, cage dust was rubbed on the apparatus. The results from this initial trial were omitted from the final analysis. The tests were performed in the order listed below with a spacing of 24 to 48 hours between tests.

**Open Field:** The rectangular arena (50 X 50cm) was cleaned, and using the ethovision software it was divided into center, border and wall zones. Animals were placed in the center of the open field arena to allow them to explore the open field area for 20 mins. The activity of the animal was videographed using Noldus Ethovision XT software. The EthoVision software automatically records the animal's movement, tracking the time spent in the center versus the border zones, as well as the total distance moved, and the speed of movement. The collected data were analyzed to determine the animal's preference for the center or border of the arena. Less time spent in the center and more in the border indicates higher anxiety-like behavior. Mean s.e.m. Statistical analyses involved two-way ANOVA with Tukey's multiple comparison correction, utilizing GraphPad Prism software (version 9). n = 16 (WT control), 33 (*Tfap2d* Het), 29(*Tfap2d* KO), 14 (WT control), 14 (Tfap*2d* cHet) and 15 (*Tfap2d* cKO).

**O-maze:** The elevated O-maze is clean and positioned in a quiet room and the EthoVision system was calibrated to recognize the different sections of the maze (open arms vs. closed arms). Animals were gently placed in the center of the closed arm of O-maze, and allowed to explore freely for 10 mins. The EthoVision software tracked the animal's movements, specifically recording entries into and the time spent in the open and closed arms. The data were analyzed to assess the animal's preference for closed arms over the open arms, which indicates anxiety-like behavior. The frequency of entries and the duration spent in each arm type are key metrics for this analysis. Statistical analyses involved two-way ANOVA with Tukey's multiple comparison correction, utilizing GraphPad Prism software (version 9). n = 14 (WT control), 38 (*Tfap2d* Het), 27 (*Tfap2d* KO), 19 (WT control), 13 (Tfap*2d* cHet) and 16 (*Tfap2d* cKO).

**Light dark box:** The assessment of anxiety levels using the light/dark box test was conducted in accordance with the methodology detailed by Bourin *et al* [65]. In summary, the experimental setup involved a dual-compartment apparatus, each compartment made of opaque Plexiglas and measuring 18 cm in length, 10 cm in width, and 13 cm in height. The light compartment was illuminated by a 100 W lamp positioned overhead, with the light diffused through a transparent Plexiglas lid. Mice were granted access between the two compartments via a small aperture connecting them. Mice were initially placed within the illuminated section. Observations commenced from the moment a mouse first entered the dark compartment, continuing for a duration of 5 minutes. The amount of time the animal spent in the light box was noted. The animals were scored blindly without information about their genotypes. The primary behavioral metrics quantified included the duration of time spent within the light compartment, measured in seconds, and the frequency of compartment transitions. Data are presented as mean ± s.e.m. Statistical analyses involved one-way repeated measures ANOVA with Tukey's multiple comparison correction, utilizing GraphPad Prism software (version 9). n = 10 (WT control), 10 (*Tfap2d* Het), and 7 (*Tfap2d* KO).

**Tail suspension:** Mice were suspended by the tail with a paper clip attached with adhesive tape about 5 mm from the end of the tail. Time spent immobile was recorded over the duration of 6 min. The video was recorded during the test. After completion of the test, mice were returned to a holding cage until all cage-mates were tested. After completion of the experiment, all mice were returned to the home cage and transferred back to the holding room. The videos were scored blindly by an independent person for the time animals were mobile and immobility time (sec) was calculated. Data are presented as mean ± s.e.m. Statistical analyses involved one-way repeated measures ANOVA with Tukey's multiple comparison correction, utilizing GraphPad Prism software (version 9). n = 18 (WT control), 24 (*Tfap2d* Het), and 13 (*Tfap2d* KO).

**Forced swim:** Animals were gently introduced into transparent 4-liter glass beakers filled with 2.5 liters of water at ambient temperature. Each subject was gently introduced into the water, ensuring their ability to maintain their nostrils above the surface for breathing. On day 1 of the experimental procedure, animals were exposed for a duration of 15 minutes. Subsequently, on the following day, the immersion time was reduced to a series of 5-minutes. The videos of the test were recorded and were scored blindly by an independent person for the time animals were mobile and immobility time (sec) was calculated. Immobility was characterized by a state of passive floating, wherein the subject performed minimal movements necessary to remain afloat. Data are presented as mean ± s.e.m. Statistical analyses involved one-way repeated measures ANOVA with Tukey's multiple comparison correction, utilizing GraphPad Prism software (version 9). n = 20 (WT control), 23 (*Tfap2d* Het) and 13 (*Tfap2d* KO).

**Fear conditioning**: The experimental cohort comprised of male and female littermates' siblings aged between PD 120 - 180. The method used for contextual and cued fear conditioning was outlined by [66,67], with some modifications. The experiment consisted of 4 parts: habituation, training, contextual memory assessment, and cued memory evaluation. During the habituation phase, mice were introduced to a dimly lit enclosure composed of dark plastic with a cardboard floor, incorporating sawdust from their original housing to establish Context A, without the administration of shocks or cues. For the training phase, the subjects were relocated to a differently structured chamber equipped with illuminated stainless steel grids capable of delivering electric shocks, designated as Context B, and infused with a cinnamon scent beneath the grids to serve as a unique olfactory stimulus. Following a 160-second acclimation period, an auditory cue (75 decibels, 2.8 kHz) was emitted for 20 seconds, with the concluding 2 seconds overlapping with an electric shock (0.5mAmp). The mice experienced six instances of tone-shock associations, interspersed with 40-second intervals, and concluded with a 60-second relaxation phase before being returned to their original cages. The subsequent day involved testing for contextual memory in the same chamber (Context B) but with the introduction of a vanilla scent beneath the grids to differentiate from the training phase's cinnamon odor, thereby focusing on contextual rather than olfactory memory recall. Cued memory was assessed in the original Context A setup, employing the same auditory cues as during training but omitting the shocks. All sessions were video recorded, and the mice's freezing behavior—defined as the absence of movement for each 3-second video frame—was analyzed. No significant variances in body weight or shock response were noted. Data are presented as mean ± s.e.m. Statistical analyses involved two-way ANOVA for the training and cued memory test and multiple comparisons with Tukey's correction, utilizing GraphPad Prism software (version 9). For conditioned memory test, one way ANOVA with Tukey's correction was applied. n = 27 (WT control), 38 (*Tfap2d* Het), 26 (*Tfap2d* KO), 16 (WT control), 12 (Tfap*2d* cHet) and 16 (*Tfap2d* cKO).

#### FOS Immunostaining and Registration to the Mouse Brain Common Coordinate Framework

To determine the activity of different brain regions after cued memory test, we performed a whole mount FOS staining using the LifeCanvas Technologies. Animals were euthanized 60-90 mins after the cued memory test using isoflourane followed by transcardial perfusion with ice-cold 1X PBS with 10 U/mL heparin until the fluid ran clear, followed by ice-cold 4% PFA. The extracted brains were incubated in 4% PFA solution at 4°C for 24hrs with gentle shaking. The samples were cleared and stained by LifeCanvas

Technologies using their protocol [68]. Samples were cleared using a SHEILD OFF solution for 3 days at 4°C, followed by incubation in the SHIELD ON for 24hrs at 37°C with light shaking. Samples were then processed for delipidation and staining using SmartBatch+. The brains after delipidation were blocked with donkey serum or 2 days, followed by primary antibody incubation for 1-3 days and secondary antibody for incubation 6h. After each primary and secondary antibody incubations, stringent washes and PFA fixation was performed. The brains were imaged using SmartSPIM at 4 μm z-step and 1.8 μm xy pixel size. After imaging, the samples underwent automated atlas registration to the Allen Brain Atlas via LifeCanvas Technologies using rigid, affine, and b-spline warping algorithms provided by SimpleElastix, for both propidium iodide (PI) and FOS (647) channels. For the KO brains where the BLC shape was altered, the protocol was modified and manually registered for atlas registration. For cell detection, custom cell detection networks developed by LifeCanvas Technologies, employing a two-step approach with a U-Net architecture for candidate detection and a 3D ResNet for classification, and were trained on hand-tagged 3D ROIs to accurately identify cell locations. The detected cells were then mapped onto the Allen Brain Atlas for quantitative analysis. Colocalization analysis determined cell coexpression of FOS and PI by measuring the euclidean distance between cells in different channels, with coexpressed cells within a 3-voxel distance threshold being counted for each atlas-defined brain region. The output generated would provide the number of FOS positive cells per $mm^3$ in each region labelled by the allen brain atlas (ABA).

**Functional Network Construction**

We developed a comprehensive analytical framework to examine the relationship between various regions functionally active after cued memory test based on the FOS expression[69,70]. The analysis was performed using Python, with key libraries including Pandas for data manipulation, Seaborn and Matplotlib for visualization, NetworkX for network analysis, and SciPy for statistical tests. We calculated Pearson correlation coefficients to assess the linear relationship between each pair of subregions [43,70] . To evaluate the statistical significance of these correlations, p-values were computed. Heatmaps were generated to visualize the correlation matrix amongst brain regions associated with threat responding, with annotations indicating the significance levels. The upper triangle of the matrix was masked to prevent redundancy, given the symmetric nature of correlation matrices. A diverging color palette from blue (negative correlation) to red (positive correlation) was applied for visual clarity. Network graphs were created to visualize the connectivity between subregions based on their correlation coefficients. Edges were drawn for correlations surpassing a defined cutoff of 0.85, emphasizing stronger relationships. Edge colors transitioned from blue to red, representing the spectrum of correlation strengths.

### Extended Data References


51 Schwab, M. H. *et al.* Neuronal basic helix-loop-helix proteins (NEX and BETA2/Neuro D) regulate terminal granule cell differentiation in the hippocampus. *J Neurosci* **20**, 3714-3724 (2000). https://doi.org/10.1523/JNEUROSCI.20-10-03714.2000

52 Guzzardo, P. M. *et al.* A small cassette enables conditional gene inactivation by CRISPR/Cas9. *Sci Rep* **7**, 16770 (2017). https://doi.org/10.1038/s41598-017-16931-z

53 Fischl, B. FreeSurfer. *Neuroimage* **62**, 774-781 (2012). https://doi.org/10.1016/j.neuroimage.2012.01.021

54 Desikan, R. S. *et al.* An automated labeling system for subdividing the human cerebral cortex on MRI scans into gyral based regions of interest. *Neuroimage* **31**, 968-980 (2006). https://doi.org/10.1016/j.neuroimage.2006.01.021

55 French, L. & Paus, T. A FreeSurfer view of the cortical transcriptome generated from the Allen Human Brain Atlas. *Front Neurosci* **9**, 323 (2015). https://doi.org/10.3389/fnins.2015.00323

56 Lim, M. M., Hammock, E. A. & Young, L. J. A method for acetylcholinesterase staining of brain sections previously processed for receptor autoradiography. *Biotech Histochem* **79**, 11-16 (2004). https://doi.org/10.1080/10520290410001671344

57 Cristancho, A. G. *et al.* Deficits in Seizure Threshold and Other Behaviors in Adult Mice without Gross Neuroanatomic Injury after Late Gestation Transient Prenatal Hypoxia. *Dev Neurosci* **44**, 246-265 (2022). https://doi.org/10.1159/000524045

58 Paxinos, G. *Atlas of the Developing Mouse Brain at E17.5, P0 and P6*. (Elsevier Science, 2007).

59 Dong, H. W. *The Allen reference atlas: A digital color brain atlas of the C57Bl/6J male mouse*. (John Wiley & Sons Inc, 2008).

60 Feng, L. *et al.* Population-averaged macaque brain atlas with high-resolution ex vivo DTI integrated into in vivo space. *Brain Struct Funct* **222**, 4131-4147 (2017). https://doi.org/10.1007/s00429-017-1463-6

61 Huang, H., Zhang, J., van Zijl, P. C. & Mori, S. Analysis of noise effects on DTI-based tractography using the brute-force and multi-ROI approach. *Magn Reson Med* **52**, 559-565 (2004). https://doi.org/10.1002/mrm.20147

62 Mori, S., Crain, B. J., Chacko, V. P. & van Zijl, P. C. Three-dimensional tracking of axonal projections in the brain by magnetic resonance imaging. *Ann Neurol* **45**, 265-269 (1999). https://doi.org/10.1002/1531-8249(199902)45:2<265::aid-ana21>3.0.co;2-3

63 Krzywinski, M. *et al.* Circos: an information aesthetic for comparative genomics. *Genome Res* **19**, 1639-1645 (2009). https://doi.org/10.1101/gr.092759.109

64 Mineur, Y. S., Ernstsen, C., Islam, A., Lefoli Maibom, K. & Picciotto, M. R. Hippocampal knockdown of alpha2 nicotinic or M1 muscarinic acetylcholine receptors in C57BL/6J male mice impairs cued fear conditioning. *Genes Brain Behav* **19**, e12677 (2020). https://doi.org/10.1111/gbb.12677

65 Bourin, M. & Hascoet, M. The mouse light/dark box test. *Eur J Pharmacol* **463**, 55-65 (2003). https://doi.org/10.1016/s0014-2999(03)01274-3

66 Mineur, Y. S., Belzung, C. & Crusio, W. E. Functional implications of decreases in neurogenesis following chronic mild stress in mice. *Neuroscience* **150**, 251-259 (2007). https://doi.org/10.1016/j.neuroscience.2007.09.045

67 Maren, S., Aharonov, G. & Fanselow, M. S. Retrograde abolition of conditional fear after excitotoxic lesions in the basolateral amygdala of rats: absence of a temporal gradient. *Behav Neurosci* **110**, 718-726 (1996). https://doi.org/10.1037//0735-7044.110.4.718

68 Park, Y. G. *et al.* Protection of tissue physicochemical properties using polyfunctional crosslinkers. *Nat Biotechnol* (2018). https://doi.org/10.1038/nbt.4281

69 Terstege, D. J. & Epp, J. R. Network Neuroscience Untethered: Brain-Wide Immediate Early Gene Expression for the Analysis of Functional Connectivity in Freely Behaving Animals. *Biology (Basel)* **12** (2022). https://doi.org/10.3390/biology12010034

70 Cho, J. H., Rendall, S. D. & Gray, J. M. Brain-wide maps of Fos expression during fear learning and recall. *Learn Mem* **24**, 169-181 (2017). https://doi.org/10.1101/lm.044446.116
